# Pathobiology of the autophagy-lysosomal pathway in the Huntington’s disease brain

**DOI:** 10.1101/2024.05.29.596470

**Authors:** Martin J. Berg, Veeranna, Corrinne M. Rosa, Asok Kumar, Panaiyur S. Mohan, Philip Stavrides, Deanna M. Marchionini, Dun-Sheng Yang, Ralph A. Nixon

**Author notes:** Equal contribution (Martin J. Berg, Veeranna, Corrinne M. Rosa, Asok Kumar). Lead contact: Ralph A. Nixon, MD, PhD Center for Dementia Research Nathan S. Kline Institute, 140 Old Orangeburg Road Orangeburg, NY 10962.

## Abstract

Accumulated levels of mutant huntingtin protein (mHTT) and its fragments are considered contributors to the pathogenesis of Huntington’s disease (HD). Although lowering mHTT by stimulating autophagy has been considered a possible therapeutic strategy, the role and competence of autophagy-lysosomal pathway (ALP) during HD progression in the human disease remains largely unknown. Here, we used multiplex confocal and ultrastructural immunocytochemical analyses of ALP functional markers in relation to mHTT aggresome pathology in striatum and the less affected cortex of HD brains staged from HD2 to HD4 by Vonsattel neuropathological criteria compared to controls. Immunolabeling revealed the localization of HTT/mHTT in ALP vesicular compartments labeled by autophagy-related adaptor proteins p62/SQSTM1 and ubiquitin, and cathepsin D (CTSD) as well as HTT-positive inclusions. Although comparatively normal at HD2, neurons at later HD stages exhibited progressive enlargement and clustering of CTSD-immunoreactive autolysosomes/lysosomes and, ultrastructurally, autophagic vacuole/lipofuscin granules accumulated progressively, more prominently in striatum than cortex. These changes were accompanied by rises in levels of HTT/mHTT and p62/SQSTM1, particularly their fragments, in striatum but not in the cortex, and by increases of LAMP1 and LAMP2 RNA and LAMP1 protein. Importantly, no blockage in autophagosome formation and autophagosome-lysosome fusion was detected, thus pinpointing autophagy substrate clearance deficits as a basis for autophagic flux declines. The findings collectively suggest that upregulated lysosomal biogenesis and preserved proteolysis maintain autophagic clearance in early-stage HD, but failure at advanced stages contributes to progressive HTT build-up and potential neurotoxicity. These findings support the prospect that ALP stimulation applied at early disease stages, when clearance machinery is fully competent, may have therapeutic benefits in HD patients.

## Introduction

Huntington’s disease (HD) is caused by a mutation in the gene encoding the huntingtin protein (HTT) resulting in expansion of the polyglutamine (polyQ) stretch on its amino-terminus (The-Huntington’s-Disease-Collaborative-Research-Group 1993, Bates, Dorsey et al. 2015, Jiang, Handley et al. 2023, Franklin, Teive et al. 2024). In the neostriatum, the brain region most vulnerable and devastated by pathology in HD, the spatiotemporal advance of atrophy dorso-ventrally, caudo-rostrally, and medio-laterally has been used to stage disease pathology severity as Grades 0-4 (HD0-HD4) (Vonsattel, Myers et al. 1985). Striatal GABA-containing medium spiny projection neurons are most susceptible to cell death, even in premanifest HD0, while other striatal neuronal populations, such as aspiny interneurons, seem more resistant to toxicity (Vonsattel 2008, Hedreen, Berretta et al. 2024). In the more advanced HD3 and HD4, pyramidal cells in Layers III, V and VI of the cerebral cortex also show cell loss (Sotrel, Paskevich et al. 1991). More recent studies have revealed that loss of Layer Va pyramidal neurons, identified as corticostriatal cells, can occur in the early stage of HD (Pressl, Mätlik et al. 2024)

Neuronal intranuclear inclusions (NII) and neuropil inclusions are present in HD brains (DiFiglia, Sapp et al. 1997) and are positive for mutant huntingtin (mHTT) and ubiquitin (Ub) (DiFiglia, Sapp et al. 1997, Becher, Kotzuk et al. 1998, Gutekunst, Li et al. 1999), suggesting that there may be a deficiency in the proteolytic machinery responsible for normally clearing these proteins, resulting in their accumulation to form the inclusions. Autophagy is generally the principal mechanism by which cells clear organelles, long-lived proteins and damaged, misfolded, or aggregated proteins that are poor substrates for the ubiquitin-proteasome system (UPS). In cell or mouse models of HD, HTT accumulates in autophagosomes (AP) and autolysosomes (AL) along with lysosomal enzyme cathepsin D (CTSD) in proportion to HTT polyQ length. This has suggested that the autophagic-lysosomal pathway (ALP) may be a major, if not the major, path for HTT proteolysis and degradation which mobilizes to clear an overload of exogenously expressed protein, particularly forms that misfold and potentially aggregate (Kegel, Kim et al. 2000, Ravikumar, Duden et al. 2002, Qin, Wang et al. 2003, Yamamoto, Cremona et al. 2006, Heng, Detloff et al. 2010). Persistent accumulation of mutant proteins/aggregates in the ALP could possibly reflect impaired autolysosomal proteolytic clearance and/or inadequate autophagy induction and flux of substrates through the ALP.

Studies in the past decade, conducted primarily with cell and/or animal models, have demonstrated multiple roles of HTT and mHTT in autophagy in relation to HD (Martin, Ladha et al. 2015, Croce and Yamamoto 2019, Klionsky, Petroni et al. 2021). Wild type HTT participates in normal autophagy by (1) releasing ULK1 from the inhibition of mTOR and (2) serving as a scaffold to facilitate cargo sequestration through improving the interaction of p62/SQSTM1 (p62) with ubiquitinated cargos and with LC3 (Ochaba, Lukacsovich et al. 2014, Rui, Xu et al. 2015). mHTT influences autophagy in HD settings in multiple ways. Although mHTT may activate autophagy through sequestering mTOR and therefore reduce mTOR activity (Ravikumar, Vacher et al. 2004), most reported mHTT effects on the ALP appear to be inhibitory for autophagy, impairing earlier stages of autophagy, including initiation signaling, phagophore nucleation and cargo recognition/AP formation. The related mechanisms involve: binding to Rheb and promoting mTOR signaling (Pryor, Biagioli et al. 2014); interfering with ULK1 activities leading to impairment of the Beclin1-PIK3C3/VPS34 and ATG14 complex (Wold, Lim et al. 2016); impairing autophagosomal cargo recognition (Martinez-Vicente, Talloczy et al. 2010, Rui, Xu et al. 2015); and interfering with the interaction between Ataxin 3 and Beclin-1, resulting in Beclin-1 degradation by the UPS (Ashkenazi, Bento et al. 2017). Additional effects of mHTT on the endolysosomal system include inducing extensive endosomal tubulation (Kegel, Kim et al. 2000), reducing exocytosis and promoting AL accumulation (Zhou, Peskett et al. 2021), and decreasing transport of late autophagic structures from the neurites to the soma (Pircs, Drouin-Ouellet et al. 2022).

It should be noted, however, that most of the above findings have been obtained from cell and/or mouse models of HD, including very recent studies using induced neurons through reprogramming human fibroblasts (Oh, Lee et al. 2022, Pircs, Drouin-Ouellet et al. 2022). Remarkably, information about potential alterations of the ALP in the human HD brain is very limited in the literature. For example, early studies show that activities of several lysosomal enzymes (β-glucuronidase, α-glucosidase, dipeptidyl aminopeptidase II and cathepsin H) are altered in brains of patients with HD (Cross, Crow et al. 1986, Mantle, Falkous et al. 1995). Fragmentary information on early neuropathological characterization of HD brain has suggested an association of HTT with endo-lysosomal compartments such as multivesicular bodies (Sapp, Schwarz et al. 1997), and an increased frequency of dystrophic neurites (Jackson, Gentleman et al. 1995, Maat-Schieman, Dorsman et al. 1999) although the extent and the specific nature and composition of dystrophic neurites in HD brain are little known.

Thus, in the HD brain, it is yet unknown whether autophagy induction is stimulated or suppressed in neurons at any stage of the disease. It is also not known whether autophagy clearance steps, such as AP-lysosome (LY) fusion and autolysosomal proteolytic function are competent. Such information, particularly defining the site(s) at which autophagy may be disrupted, is crucial to designing possible interventions for HD based on autophagy modulation. Therefore, in this study, we aimed to characterize the status of the ALP in relation to disease progression in human brain samples, employing ultrastructural, immunohistochemical, and molecular analyses for the caudate nucleus of the striatum (STR) and the prefrontal cortex (CTX) from control cases and HD2-HD4 patients. Importantly, for these analyses, we accessed a postmortem HD brain with exceptionally well-preserved ultrastructural preservation enabling detailed neuropathological and immunochemical analyses of autophagy-related alterations. To our knowledge, this study is unique in terms of the relatively large number of HD cases used (**Table 1**), the level of ultrastructural analysis, the range of autophagy related processes analyzed, and the value of human brain-derived information regarding this disease.

**Table 1.**
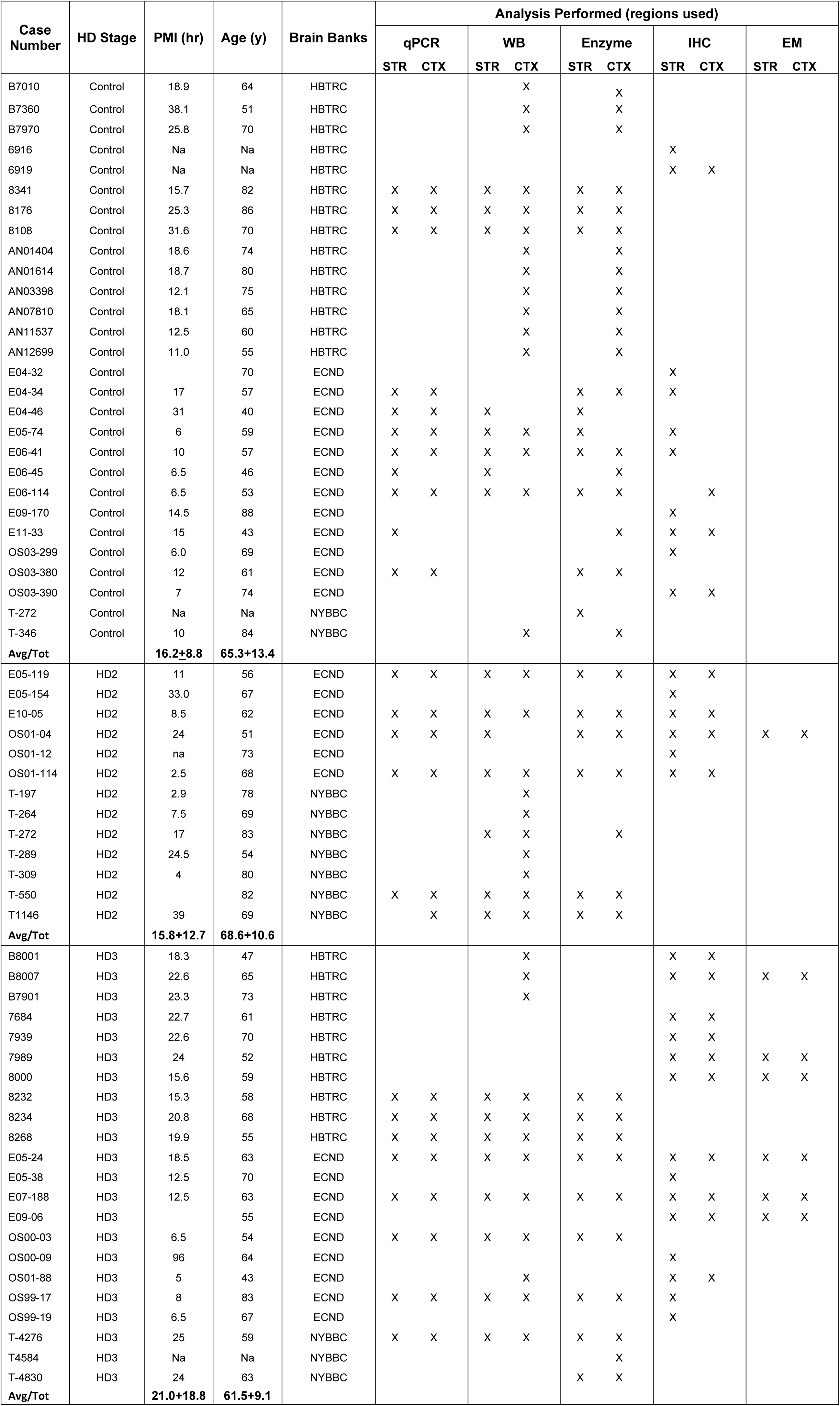
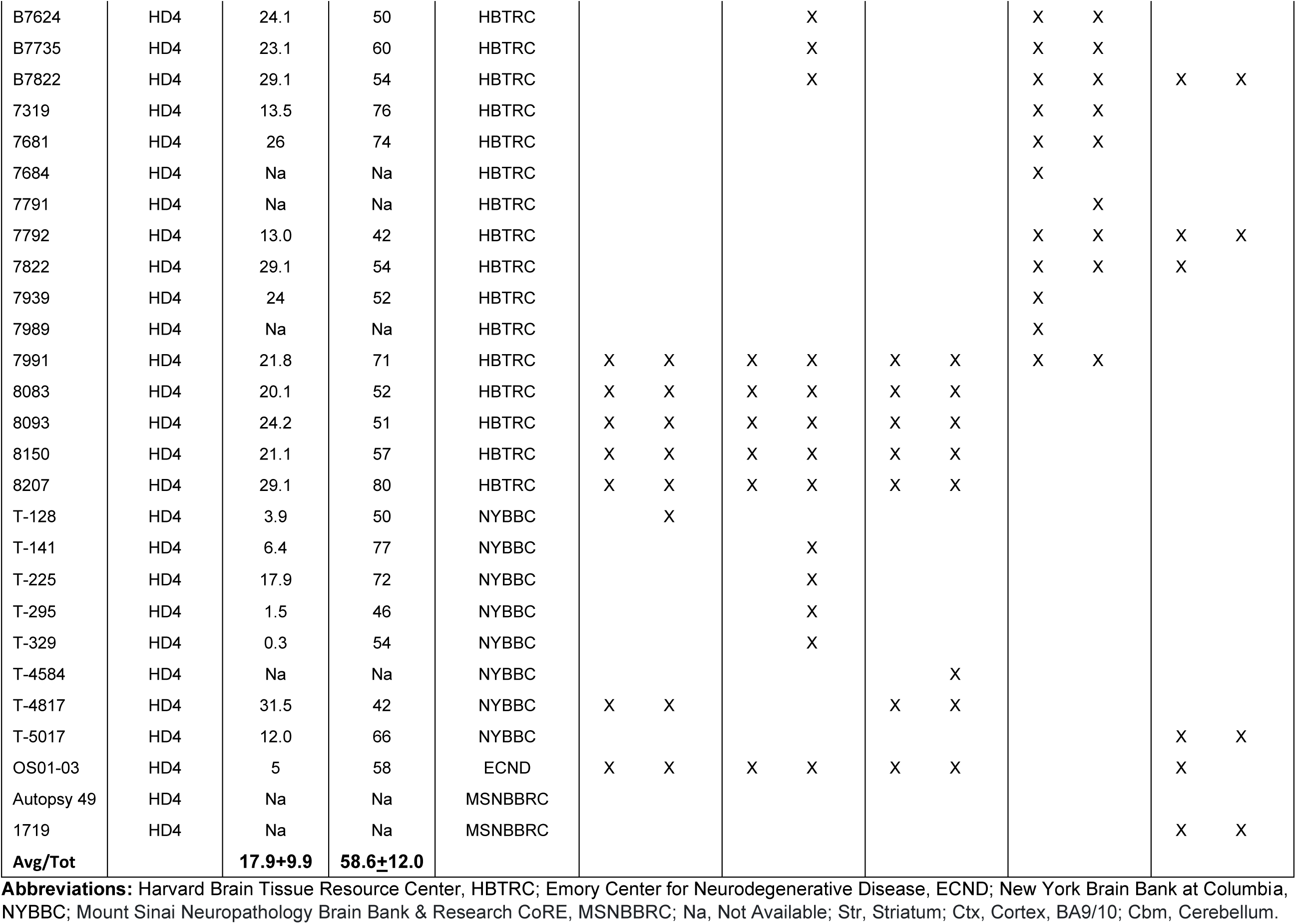
Demographics of human brain samples utilized for qPCR for mRNA analysis (qPCR), western blotting (WB), cathepsin enzymatic activity assay (Enzyme), immunohistochemistry (IHC) and electronic microscopy (EM).

## Results

### HTT inclusions: types and close relationships with autophagy adaptor proteins Ub and p62

Consistent with previous studies reporting NIIs in human HD brains (DiFiglia, Sapp et al. 1997), our ultrastructural analysis of affected neurons in the STR and CTX revealed nuclei containing single discrete ovoid or irregular shaped NIIs, 1-4 µm in diameter, composed of relatively uniform meshwork of granular or short fibrous elements (Fig. 1A1, arrowheads), which were not detected in control human brains (Fig. S1a). At the light microscope level, NIIs were readily detected by antibodies directed against HTT (mEM48) or autophagy adaptor proteins including Ub and p62 (Fig. 1A2). The extent to which nuclei were filled by mHTT, Ub or p62 immunoreactivity (IR) varied, ranging from only a small punctate, or the rimming of the outer surface of the nuclear envelope, to progressive labeling of the entire nucleus (Fig. S1b – using Ub as an example). Double immunofluorescence labeling detected a high degree of colocalization of nuclear p62-IR and Ub-IR (Fig. 1A3).

**Fig. 1.**
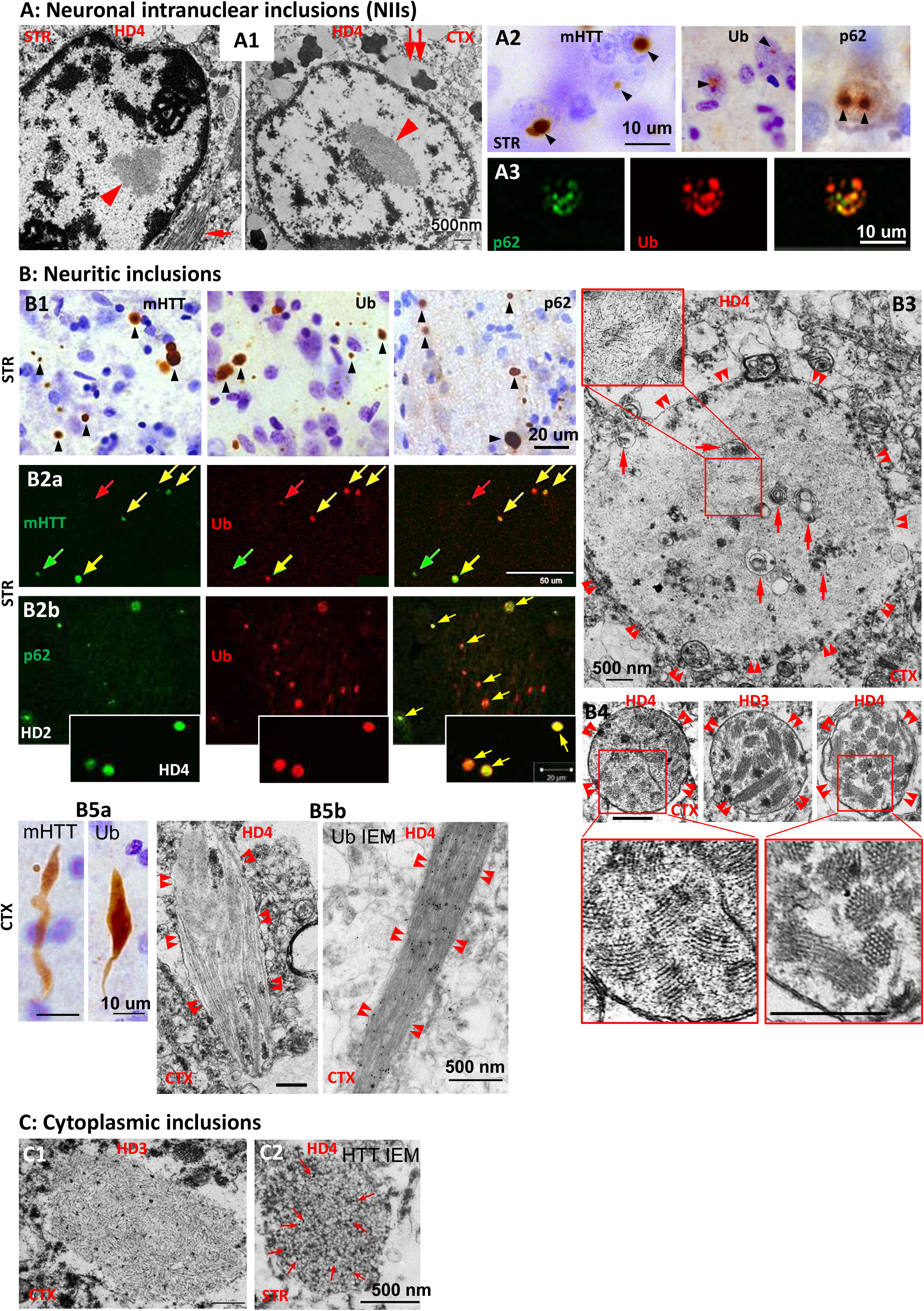
HTT inclusions: types, distributions and close relationship with autophagy adaptors Ub and p62. (A) Representative micrographs taken from the STR and/or CTX (as depicted on the individual panels) of HD brains for demonstrating NIIs. (A1) EM images depicting the NII (arrowhead). Bar = 500 nm. (A2) LM images showing that NIIs (arrowheads) are detected by antibodies to mHTT (mEM48), pan-Ub or p62. Bar = 10 μm. (A3) Confocal images from brain sections double-labeled with anti-p62 and -Ub antibodies depicting a high degree of colocalization of the two markers within NIIs. Bar = 10 μm. (B) Representative micrographs taken from the STR and/or CTX (as depicted on the individual panels) of HD brains for demonstrating neuritic inclusions. (B1) LM images showing neuritic inclusions (arrowheads) in the neuropil detected by antibodies to mHTT (mEM48), pan-Ub or p62. Bar = 20 μm. (B2) Confocal images from brain sections double-labeled with anti-mHTT (mEM48) and -pan-Ub (B2a), or anti-p62 and -pan-Ub (B2b) antibodies demonstrating labeled neuritic inclusions (arrows) in the neuropil. Yellow arrows depict double-labeled inclusions indicating colocalization, while red and green arrows point to singly labeled inclusions without colocalization. Bar = 50 μm (B2a) and = 20 μm (B2b). (B3) A representative EM image for the most common type (see text) of neuritic inclusions (double arrowheads depict the boundary of the neurite), usually 1-8 μm in diameter, filled with short, fine fibrous elements (Inset). Arrows indicate AVs within the inclusions. Bar = 500 nm. (B4) Representative EM images for the less common type (see text) of neuritic inclusions (circled by double arrowheads), usually 0.5-2 μm in diameter, characterized by a fingerprint feature, appearing to be composed of bundles of filaments/microtubules. Bar = 500 nm. (B5) Representative LM (B5a) and EM images (B5b) for the least common type (see text) of neuritic inclusions (surrounded by double arrowheads). (B5a) shows the elongated feature of the inclusions revealed by mHTT (mEM48) or pan-Ub antibodies. Bars = 10 μm. (B5b) depicts the filaments/microtubules revealed by either conventional EM or IEM with an anti-pan-Ub antibody, indicating specific immunogold labeling on the filaments. Bars = 500 nm. (C) Representative EM micrographs taken from HD brains for demonstrating cytoplasmic inclusions, which exhibit as either accumulation of fibrous filaments devoid of a limiting membrane (C1), or accumulation of very small (< 30 nm in diameter) clear vesicles which are positive for mHTT (mEM48) as shown by labeling of immunogolds (red arrows) (C2). High resolution images for (C) are presented as Fig. S1d.

Another major type of inclusion was neuritic inclusions randomly distributing in the neuropil. They were also readily detected by anti-mHTT, -Ub and -p62 antibodies in the STR (Fig. 1B1, B2) and the CTX (Fig. S1c). The majority of neuritic inclusions exhibited spherical or oval shapes, with hugely varied sizes (0.5-8 µm) (Fig. 1B1, B2; Fig. S1c). High levels of colocalization were observed between mHTT-IR and Ub-IR (Fig. 1B2a) or p62-IR and Ub-IR (Fig. 1B2b), implying colocalization of the three proteins within the inclusions, which was seen in our study in the brain of a knock-in HD mouse model Q175 (Stavrides, Goulbourne et al. 2024 bioRxiv). The extent of overlapped area of the two colors from p62-IR and Ub-IR appeared to increase with disease progression (Fig. 1B2b, HD2 vs HD4), suggesting that p62 and Ub proteins, and mHTT as well, each can aggregate independently earlier and then further develop to form composite structures. Besides these spherical/ovoid neuritic inclusions, there were elongated types of modestly enlarged neurites (Fig. 1B5a) with much lower frequency – for example, they were hardly found even in images at low magnification (Fig. 1B1; Fig. S1c).

The neuritic inclusions in the neuropil appeared to fall into 3 forms ultrastructurally. (1) Most common were 1-8 µm spherical inclusions that contained no evident limiting membrane and often occupied the entire cross-sectional area of a given neurite, leaving only a narrow surround of cytoplasm between the inclusion and the plasma membrane of the neurite (Fig. 1B3). The internal structure of these inclusions, like inclusions in the nucleus, consisted of short, fine fibrous elements (Fig. 1B3, Inset). Autophagic vacuoles (AVs) or other small membranous vesicles were sometimes trapped within these structures (Fig. 1B3, arrows). (2) The 2^nd^ type of inclusion in the neuropil, which were less common than the above inclusions, but still frequently seen, were 0.5-2.0 µm membrane-bound spherical structures containing multiple smaller fingerprint profiles, most of which appeared to be composed of well-arranged bundles of filaments or microtubules (Fig 1B4). These two forms of inclusions may correspond to the spherical /ovoid type found at the light microscopic level (Fig. 1B1, B2). (3) The least common inclusions in the neuropil were elongated or comet-shaped structures (Fig. 1B5b), most likely corresponding to the aforementioned elongated type under light microscope (Fig. 1B5a), which were partially or fully occupied by microtubule-like elements (Fig. 1B5b, left) or fibrillar bundles which were positive for Ub as detected by Immuno-Gold EM (IEM) (Fig. 1B5b, right).

In addition to the NIIs and the neuritic inclusions, there were cytoplasmic inclusions, which, ultrastructurally, could be identified as 3 forms. (1) one or several large (∼3 µm) ovoid or spherical structures devoid of limiting membrane and consisting of mainly a meshwork of short, thin fibrous elements intermixed with small numbers of membranous elements (Fig. 1C1; Fig. S1d). These inclusions resembled the similar size fibrous structures seen in the neuropil (Fig. 1B3). (2) 2-5 µm membrane-bound profiles containing collections of very small clear vesicles (<30 nm diameter) (Fig. 1C2; Fig. S1d). (3) Fiber bundles similar to those found in the neuropil (e.g., Fig. 1B5b) were occasionally observed perinuclearly within the cytoplasm (Fig. 1A1, STR, arrow).

### Immunoblotting analyses of HTT proteins and adaptor proteins p62, TRAF6 and Ub

The above morphological observations suggest close spatial relationships among mHTT and the adaptor proteins Ub and p62. Both proteins are involved in UPS and ALP proteostasis under physiological conditions. In the pathological situation of abundant mHTT aggregates, however, ALP may become more influential as these lesions are poor substrates for the UPS. We further investigated such relationships biochemically by immunoblot analyses of these proteins in the STR and the CTX from HD patients (Vonsattel neuropathological stages HD2-HD4) and corresponding controls (Fig. 2). It should be noted that a large number of striatal and cortical samples were derived from the same patients, thus lending a great deal of confidence in comparing and contrasting these cohorts (see **Table 1**, Demographics). That said, we notably observed regionally specific patterns in the levels of HTT aggregates, intact HTT protein and HTT fragments.

**Fig. 2.**
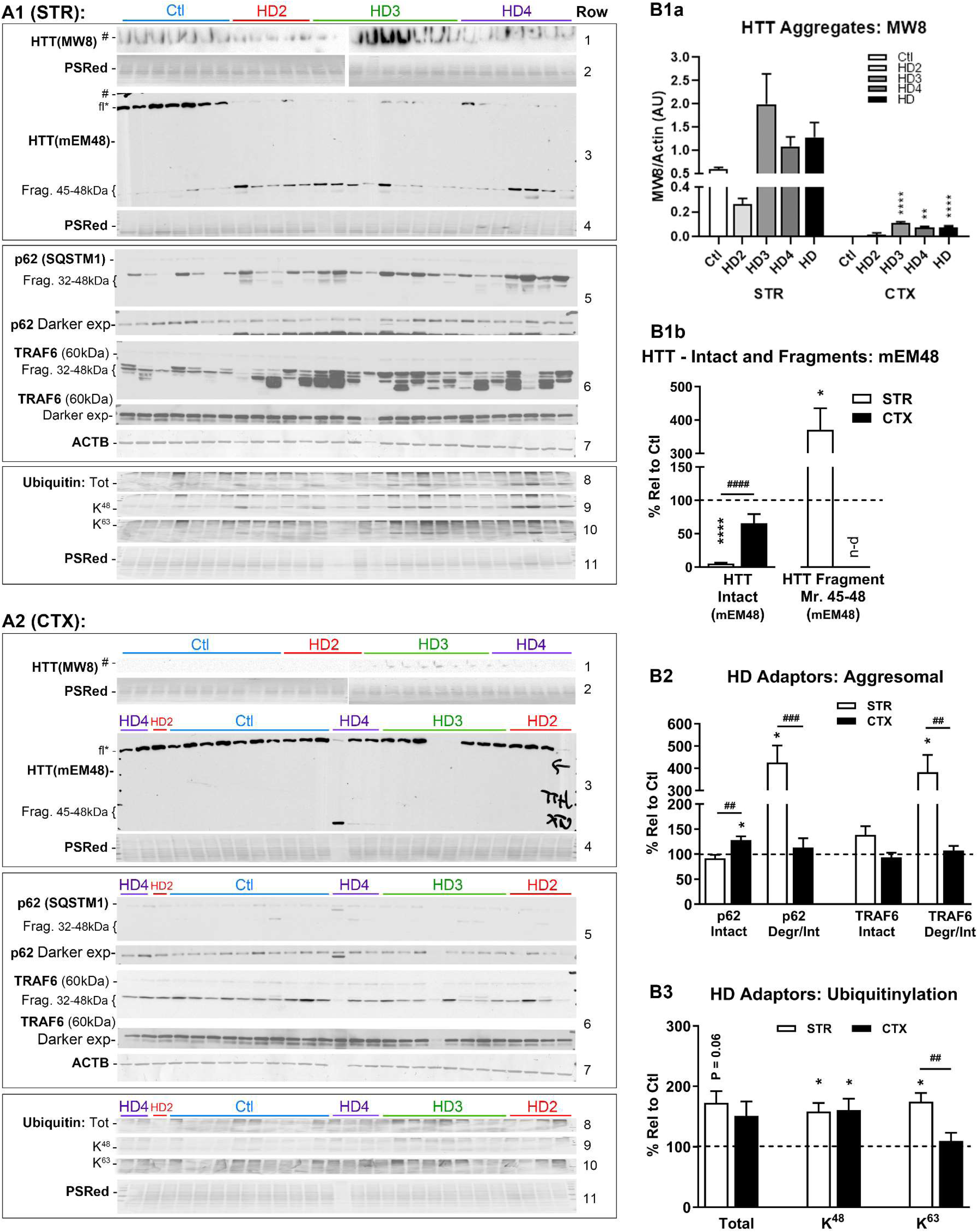
Protein levels of HTT and autophagy adaptor Ub, p62 and TRAF6 in the STR and CTX. (A) Western blots of samples from the STR (A1) and CTX (A2) for assessing the levels of HTT aggregates (with antibody MW8), HTT intact and/or fragmented species (with antibody mEM48), as well as UPS/autophagy adaptor proteins including p62, TRAF6 and Ub. All tissue samples were run at the same time in wide-format gels, electroblotted onto same membrane for immunodetection by ECL (Representative Ponceau S Red-stained blots or ACTB blots detected by colorimetric assay using DAB are shown). **Color bars above the blots denote HD staging (Blue = Control/Ctl; Red = HD2; Green = HD3; Purple = HD4)**. (B) Bar graphs showing the quantitative results for HTT aggregates (B1a), intact and fragmented HTT species (B1b), adaptor protein/aggresomal markers (B2), and adaptor protein/ubiquitination species (B3). Except B1a where data from each HD stage are shown to better demonstrate the difference of signals between the STR and CTX, each bar in other graphs (i.e. B1b, B2, B3) represents a single result of HD data pooled from all HD2, HD3 and HD4 samples and expressed as % of Ctl (set as 100% depicted by the dashed line) ± SEM. n-d: not detectable. Significant differences between the Ctl and the HD case of each brain region, or between the STR and the CTX of HD case were analyzed by two-tailed Student’s t-test. * signs: comparisons with the Ctl; # signs: comparisons between STR and CTX. * or # P<0.05, ** or ## P<0.01, *** or ### P<0.001, **** or #### P<0.0001. For STR, n = 7 control and 21 HD cases, for CTX, n = 10 control and 18 HD cases. Please note that the images of blots for ubiquitination species were compressed in the vertical direction to be 15% relative to the original height, but quantitation was done on the original blots. PSRed = Ponceau S Red staining.

First, to evaluate mHTT aggregation, we performed immunoblotting with MW8, an antibody targeting the C-terminus of exon 1 protein located in the N-terminus of HTT and capable of detecting the aggregated form of HTT and the exon 1 protein (Ko, Ou et al. 2001, Baldo, Paganetti et al. 2012), which showed HTT immunoreactivity rimming sample wells within the stacking gel region that, in HD striatal samples, increased and peaked in HD3 (Fig. 2A1, Row 1; Fig. 2B1a, left))(see Fig. S2 for the uncropped full-size blots). Only trace levels were detected in cortical samples; however, levels were greatest in HD3 as well. (Fig. 2A2, Row 1; Fig. 2B1a, right). Overall, the levels were roughly 20-fold higher in the STR versus the CTX, suggesting that aggregation occurs in a regionally specific manner.

Second, the levels of intact HTT (i.e., full-length HTT) were greatly reduced in the STR of HD cases (as early as HD2) compared to controls when detected by another N-terminal antibody, mEM48 (Gutekunst, Li et al. 1999) (Fig. 2A1, Row 3, fl (full-length)*; Fig. 2B1b left), while the difference in the levels of this HTT species in the CTX between the control and HD cases was only marginal (Fig. 2A2, Row 3, fl*; Fig. 2B1b left). The decline of the intact HTT in the STR of HD stages was accompanied by a marked accumulation of 45-48 kDa N-terminal fragments of HTT (Fig. 2A1, Row 3; Fig. 2B1b right), while such fragments were absent in the CTX (Fig. 2A2 Row 3; Fig. 2B1b right) (see Discussion).

Immunoblotting analyses of p62 and TRAF6, two adaptor proteins known to interact with each other, revealed that these two proteins also generated proteolytic fragments (32-48 kDa) which, interestingly, increased also selectively in the HD STR at a level of nearly 4-fold over the controls (Fig. 2A1, Rows 5, 6; Fig. 2B2), similar to the selective accumulation of mHTT fragments in the HD STR described above. By contrast, there were no differences in the levels of such fragments between HD and control cases in the CTX (Fig. 2A2, Rows 5, 6; Fig. 2B2). In addition, analysis of ubiquitination revealed a significant increase in K^48^ and K^63^ ubiquitination and a strong trend of elevation in Total ubiquitination (P = 0.06) in the STR and a significant increase only in K^48^ ubiquitination in the CTX (Fig. 2A1 and 2A2, Rows 8-10; Fig. 2B3). Together, the generation/accumulation of proteolytic fragments of mHTT, p62 and TRAF6, and the increase in ubiquitination, in the STR appear to be disease-related and brain region-selective.

### ALP pathology develops at the late stages of the disease as revealed by CTSD immunolabeling

We subsequently assessed the status of the ALP in HD brains, compared to control brains, with immunohistochemistry (IHC) of CTSD, which captures the status of both LY and AL collectively. AL can be distinguished from LY by residual presence of LC3 or another autophagy-specific substrate. Anti-CTSD antibody labeled many neurons in multiple layers of the CTX in control cases with small positive puncta (visible in some neurons even at a low magnification) (Fig. 3A). Brain sections at HD2 stage exhibited a similar CTSD labeling pattern (Fig. 3B) to that seen in the Controls, suggesting an absence of lysosomal pathology at this earlier stage of the disease progression. However, a mild abnormal staining pattern was observed in sections from HD4, exhibiting as stronger, clumping intraneuronal IR particularly at the basal poles, and an overall patchy staining pattern likely due to neuronal loss and increased staining in the neuropil (Fig. 3C). Similarly, in the STR, an apparently normal CTSD staining pattern was seen in sections from control cases even though there was some small clumping staining (Fig. 3D), and again, abnormal staining pattern was evident at HD4 stage reflected by strongly stained puncta and clumps (of puncta) within neuronal bodies or in the neuropil (Fig. 3E), which was more severe than that seen in the CTX (Fig. 3C). At a higher magnification (Fig. 3F-I), small and moderately CTSD-positive puncta, representing normal AL/LY morphology, were seen in neurons of control and HD2 cases (Fig. 3F, G). By contrast, strongly stained, grossly enlarged CTSD granules in neurons and neuropil became predominant at HD4 (Fig. 3I), indicating obvious lysosomal pathology. As expected, the STR at HD3 stage exhibited a CTSD staining pattern in between what seen in HD2 and HD4, i.e., some neurons were still normal appearing (Fig. 3H), while others did become abnormal (Fig. 3H, inset). Together, lysosomal pathology, as revealed by CTSD IHC, develops severely in the STR and mildly in the CTX of HD brains late in the disease progression, particularly at HD3-HD4 stages.

**Fig. 3.**
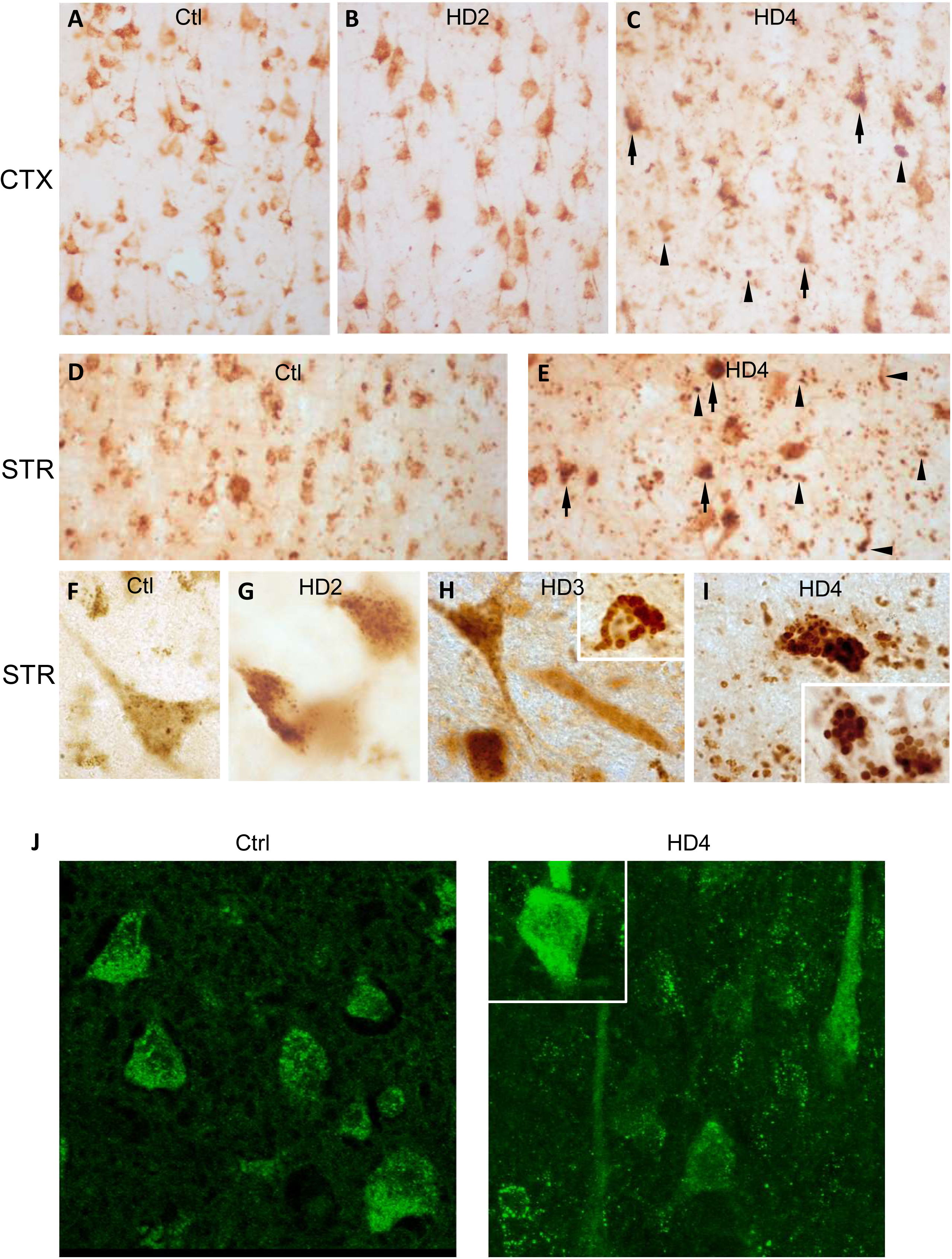
HD brains develop ALP pathology in the later stages of the disease progression, as revealed by immunostaining of CTSD. (A-I) Brain sections from control and HD cases were immunostained with an anti-CTSD antibody. Low magnification images were taken from the CTX of Control (Ctl) (A), HD2 (B) and HD4 (C) stages and from the STR of Ctl (D) and HD4 (E) depicting the normal CTSD staining pattern in the Ctl brain while abnormal pattern in the HD4 brain represented by strong and clumping IR at one pole of the neurons (arrows) and strong and increased staining in swollen neurites in the neuropil (arrowheads). (F-I) High magnification images from the STR showing small punctate CTSD granules (i.e., AL/LY) in neurons of Ctl, HD2 and HD3 (F-H), while grossly enlarged positive granules in HD3 (H, Inset) and dominantly in HD4 (I and Inset). (J) Representative confocal images from Ctl and HD4 cases labeled for LC3, showing a lack of obvious alteration in LC3 staining pattern even at the late disease stage, except that some degenerating neurons exhibited enhanced diffuse staining in the cytoplasm (Inset).

Immunolabeling with LC3, another marker for the ALP, revealed LC3 puncta in neurons in the HD STR which, even at HD4, were not evidently increased in number beyond the range seen in control brains (Fig. 3J). Isolated neurons undergoing degeneration often labeled strongly but diffusely with LC3 antibodies suggesting upregulation of LC3-I but not necessarily increased formation of LC3-II puncta (Fig. 3J, inset). In general, the numbers of LC3 puncta and size distribution did not reflect any abnormal proliferation or enlargement due to impairment in a particular step of autophagy (e.g., AP formation or AP-LY fusion).

### Association of mHTT signal with CTSD IR during disease progression

Considering the occurrence of lysosomal pathology in HD brains appearing late in the pathology, we next evaluated possible relationships of the lysosomal abnormality with the potential substrates (e.g., HTT) reaching LYs through the autophagic process using immunofluorescence. Double immunolabeling for CTSD and HTT (mEM48) in striatal neurons from control cases revealed moderately stained CTSD puncta (Fig. 4A). HD neurons revealed mHTT immunolabeling with mEM48 that was surrounded by CTSD (Fig. 4B, C, red boxes, and 4D, arrowheads). Also, at HD4, a splotchy CTSD staining pattern was evident by the fluorescent labeling (Fig. 4C), similar to that revealed by DAB staining shown above (Fig. 3C, E), and CTSD and/or HTT positive vesicles in many cells had become grossly enlarged and clustered (Fig. 4D). The relatively weak intraluminal HTT signal at HD2 (Fig. 4B) vs. the strong signal at HD4 (Fig. C, D) may imply a relatively competent lysosomal degradative function in clearing HTT in the early stage and a progressive delay in the clearance of this substrate in overtly degenerating neurons during disease progression. At HD4, HTT-positive inclusions in the neuropil were numerous and often larger (Fig. 4C, blue boxes), colocalized with (therefore seen as yellow fluorescence), or surrounded by, CTSD IR puncta, suggesting that the HTT inclusions were too large to be contained within individual ALs. These structures in the neuropil may correspond to the neuritic inclusion shown in Fig. 1B3 where AVs exist inside or surrounding a large inclusion. The lysosomal pathology in affected neurons in late-stage HD brains (Figs. 3, 4) involved the accumulation of CTSD positive vesicles, which were decorated by HTT (Fig. 4B-D) and concentrating in one pole of the neuron (Fig. 3C, 3E; Fig. 4D), a distribution pattern associated with lipofuscin granules. This was consistent with the abundance of lipofuscin granules and clusters seen ultrastructurally (see below) and was further verified by IEM where lipofuscin granules were positively labeled by 4 markers, CTSD, p62, Ub and HTT (Fig. 4E).

**Fig. 4.**
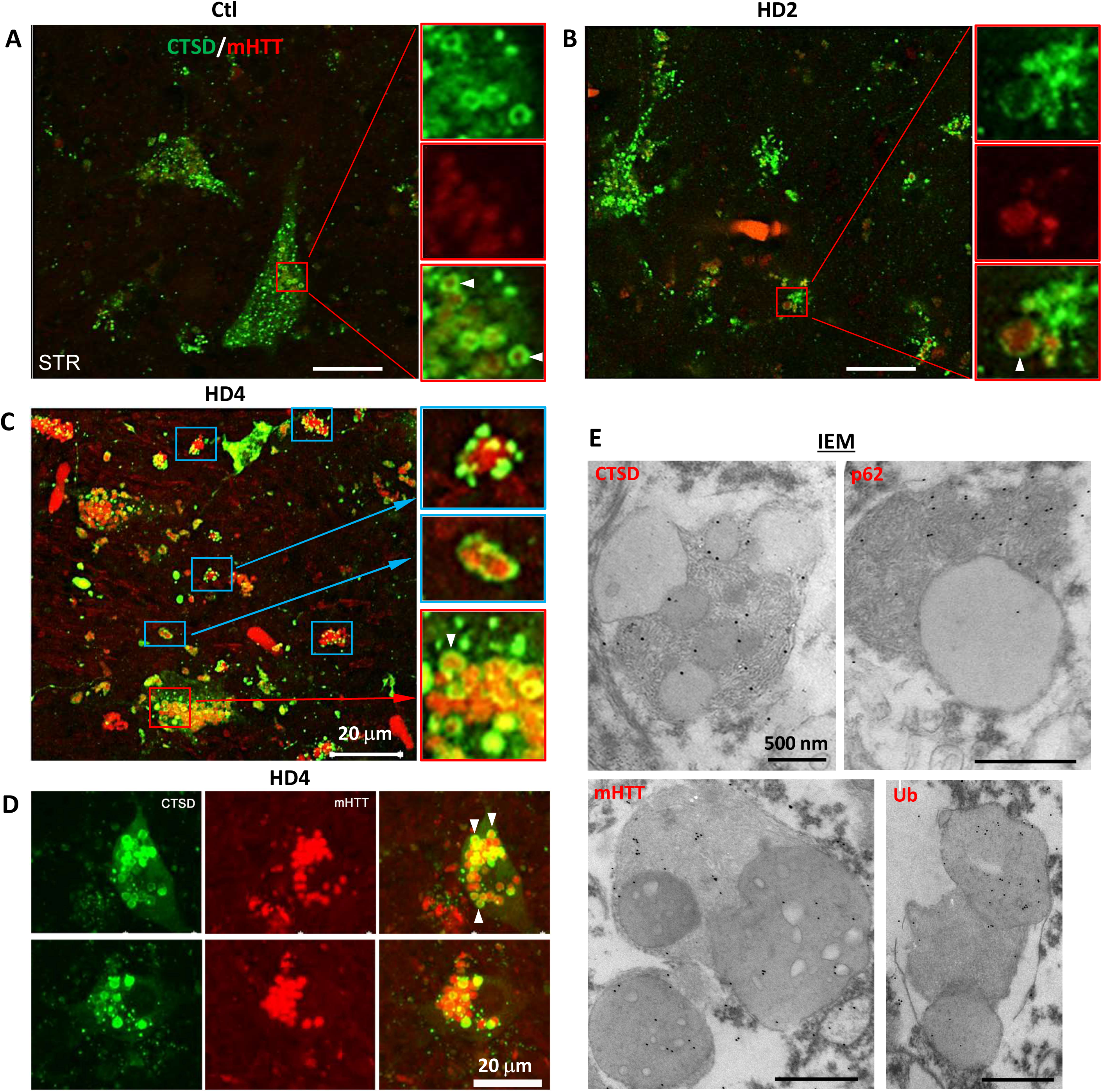
Association of mHTT with CTSD IR during disease progression. (A-D) Representative confocal images taken from the STR from control (A), HD2 (B) and HD4 (C, D) cases double labeled for mHTT (mEM48) and CTSD, depicting association/colocalization of mHTT IR in CTSD positive vesicles (AL/LY), where CTSD signal strongly decorates the rim of the lumen (arrowheads). CTSD signal also decorates mHTT-positive neuritic inclusions in the neuropil (blue boxes), in the forms of either discrete puncta (C, upper Inset) or continuous ring (C, middle Inset). (D) depicts grossly enlarged (note that D has the same magnification as A-C) and clustered vesicles at HD4 stage, positive for CTSD or mHTT or both. Bars = 20 μm. (E) Representative IEM images of HD brain samples labeled with antibodies against either CTSD, mHTT (mEM48), pan-Ub and p62, particularly demonstrating the labeling of gold particles on lipofuscin granules. Bars = 500 nm.

### Further ultrastructural analyses of cytoplasmic vesicular organelles

The above single and double immunostaining demonstrate that a dominant vesicular pathology in affected neurons of late-stage HD involves the accumulation and enlargement of CTSD positive vesicles which correspond to AL/LY and lipofuscin granules. We then extended our EM analysis to verify the incidence of these structures and to further assess autophagy related or unrelated vesicular structures in brain samples from a HD4 case, in addition to the previously described ultrastructures of cytoplasmic inclusions shown in Fig. 1C.

First, we verified that the cytoplasm of affected neurons contained a range of AVs (AP, AL, LY) (<500 nm) and abundant lipofuscin granules (which could also be referred to as pigmented ALs)(Sulzer, Mosharov et al. 2008) with either typical bipartite lipid/protein morphology (double arrows in Fig. 5A1, top inset; also see Fig. 1A1, CTX, top) or clusters of more amorphous granules with heterogeneous content of varying electron density corresponding to early forms of lipofuscin granules (Fig. 5A1, arrows; also see Fig. 1C2, right), all reflecting incomplete degradation of lysosomal substrates within. Some of these small lipofuscin granules or granule clusters were surrounded by a double membrane suggesting attempts by the cell to digest these structures by lysophagy, a process of AL/LY autophagy (Fig. 5A1, bottom inset). Second, of particular note were the collections of single membrane-limited 300-500 nm vesicles in some affected neurons, exhibiting a relatively uniform content of mainly granular and fibrous material (Fig. 5A2, single arrowheads) resembling in part the components of large fibrous cytoplasmic inclusions described above (Fig. 1C1). This population of vesicles, presumably ALs, coexisted with double membrane-limited structures, presumably APs (Fig. 5A2, double arrowheads) in the perikarya, implying no evident block in AP-LY fusion or generation of ALs. The large number of these AVs in some neurons support the concept that either the generation of APs/ALs has increased or more consistent with our other data, that clearance of ALs has decreased. Third, in dystrophic neurites (Fig. 5B1, B2), which were not frequent in the HD brain, AVs were readily identified which were present as double-membrane bound vacuoles (i.e. APs) or multilamellar bodies (MLB) containing low electron-density intralumenal content, consistent with successful sequestration and delivery of autophagy cargos along the ALP.

**Fig. 5.**
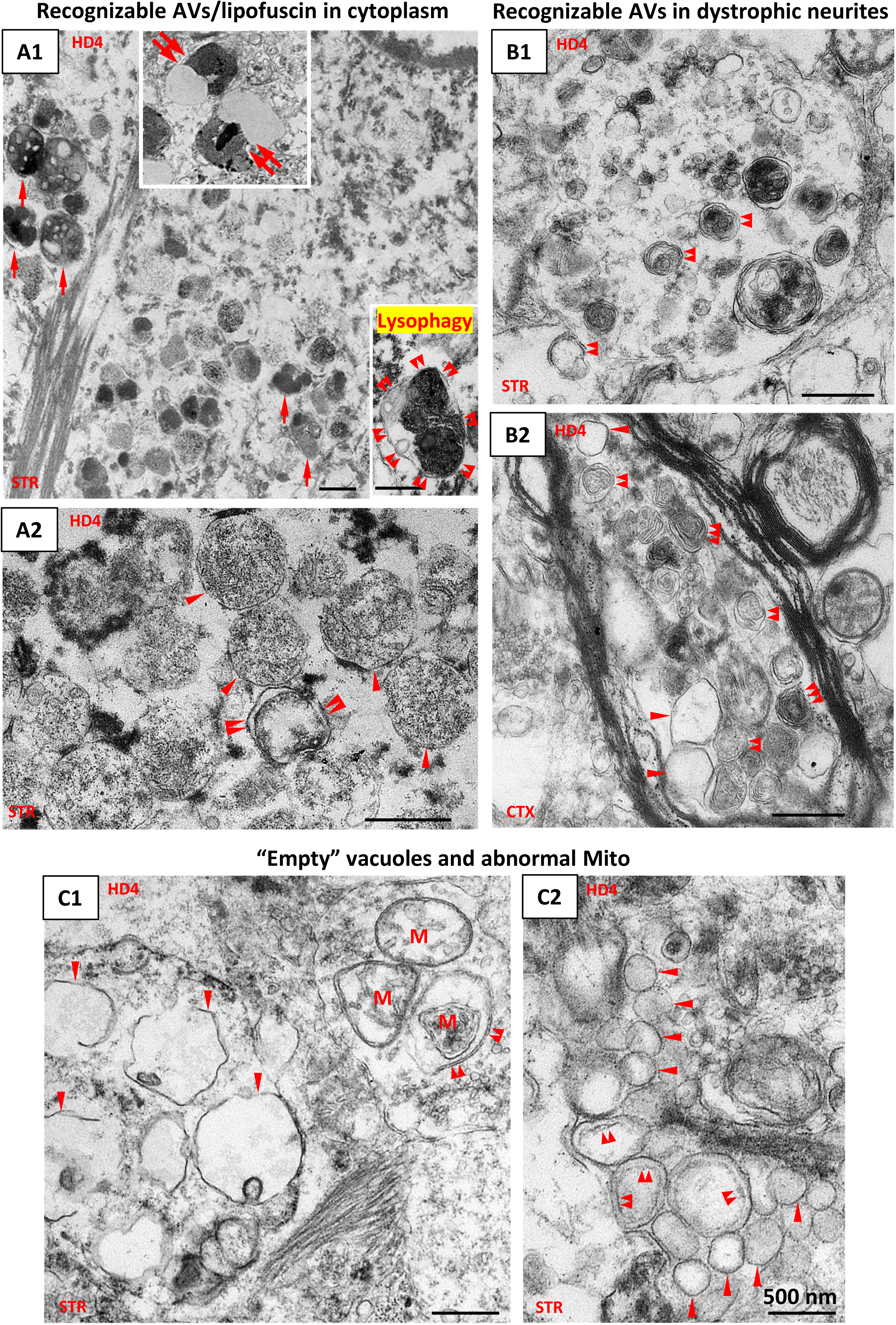
Membranous/vesicular pathologies in the later stages of HD – Accumulation of vesicles with various types of intraluminal features. Representative EM images from the STR and the CTX (as depicted on the individual panels) of a HD4 case demonstrating cytoplasmic membranous/vesicular pathologies. (A) Representative EM images depicting recognizable cytoplasmic vesicles of the ALP, including mature lipofuscin granules with bipartite protein/lipid morphology (A1, upper Inset, double arrows), early forms of lipofuscin granules (A1, single arrows), double (double arrowheads) or single (single arrowheads) membrane-limited vesicles (A2), presumably corresponding to APs and ALs respectively. The lower Inset in A1 depicts a compounded dense AL/lipofuscin granule within a double membrane sac, implying lysophagy. Bars = 500 nm. (B) Representative EM images of dystrophic neurites depicting recognizable AVs including APs with double limiting membrane (double arrowheads) and multilamellar bodies (MLB, triple arrowheads). Clear vesicles with apparent single limiting membrane within the same dystrophic neurite (B2) are indicated by single arrowheads. Bars = 500 nm. (C) Representative EM images for other vesicular structures of unidentified origins, including single membrane-limited clear vesicles of varying sizes with minimal intralumenal contents (C1, C2, single arrowheads), similar clear vesicles exhibiting apparent double membrane (C2, double arrowheads. Note that the inner membrane is faint, questioning a fixation artifact). The double arrowheads in C1 point to a double membrane vacuole containing a mitochondrion. Bars = 500 nm.

Notably, there were clusters of clear vesicles of varying sizes characterized by their minimal intralumenal content (i.e. “empty”), most of which were single membrane-limited (Fig. 5C1, C2, single arrowheads; also see Fig. 5B2), while some exhibited a double membrane appearance (Fig. 5C2, double arrowheads), raising the possibility that the latter type represents the “empty APs” due to defective cargo sequestration (Martinez-Vicente, Talloczy et al. 2010). On the other hand, however, mitochondria were seen in double membrane vesicles (Fig. 5C1, double arrowheads), implying that at least sequestration of this cargo, is unimpeded.

### Immunoblotting and qPCR of lysosomal markers suggest upregulation of lysosomal biogenesis in the STR

To survey ALP-related biochemical changes in the HD brain using immunoblotting, we analyzed protein markers of the different phases of the ALP (i.e., autophagy induction signaling, AV formation and lysosomal substrate clearance) on staged brains from HD2 - HD4 and control cases compared in two brain regions – the STR and the CTX. Notably, the majority of marker proteins for the early phases of the ALP, including p-p70S6 (as an indicator for mTOR activity), BECN1, VPS34, ATG5, ATG7 and LC3, showed no changes apparent with disease progression in both the STR and the CTX (not shown).

Next, for markers relating to lysosomal degradative functions at the later phase of the ALP, we did not observe significant alterations in the protein levels (Fig. 6A, C), and enzymatic activities (Fig. 6B, D), of lysosomal enzymes CTSD and CTSB involved in substrate clearance in the STR (Fig. 6A, B) and the CTX (Fig. 6C, D). However, an intriguing observation was a striking elevation of the levels of LAMP1 in the STR in the HD brains, appearing as early as HD2 (Fig. 6A1, A2), which was otherwise not observed in the CTX (Fig. 6B1, B2), representing a major different observation between the two brain regions and consistent with aforementioned CTSD IHC results showing severely-altered vs. mildly-altered CTSD staining patterns in the STR vs the CTX, respectively.

**Fig. 6.**
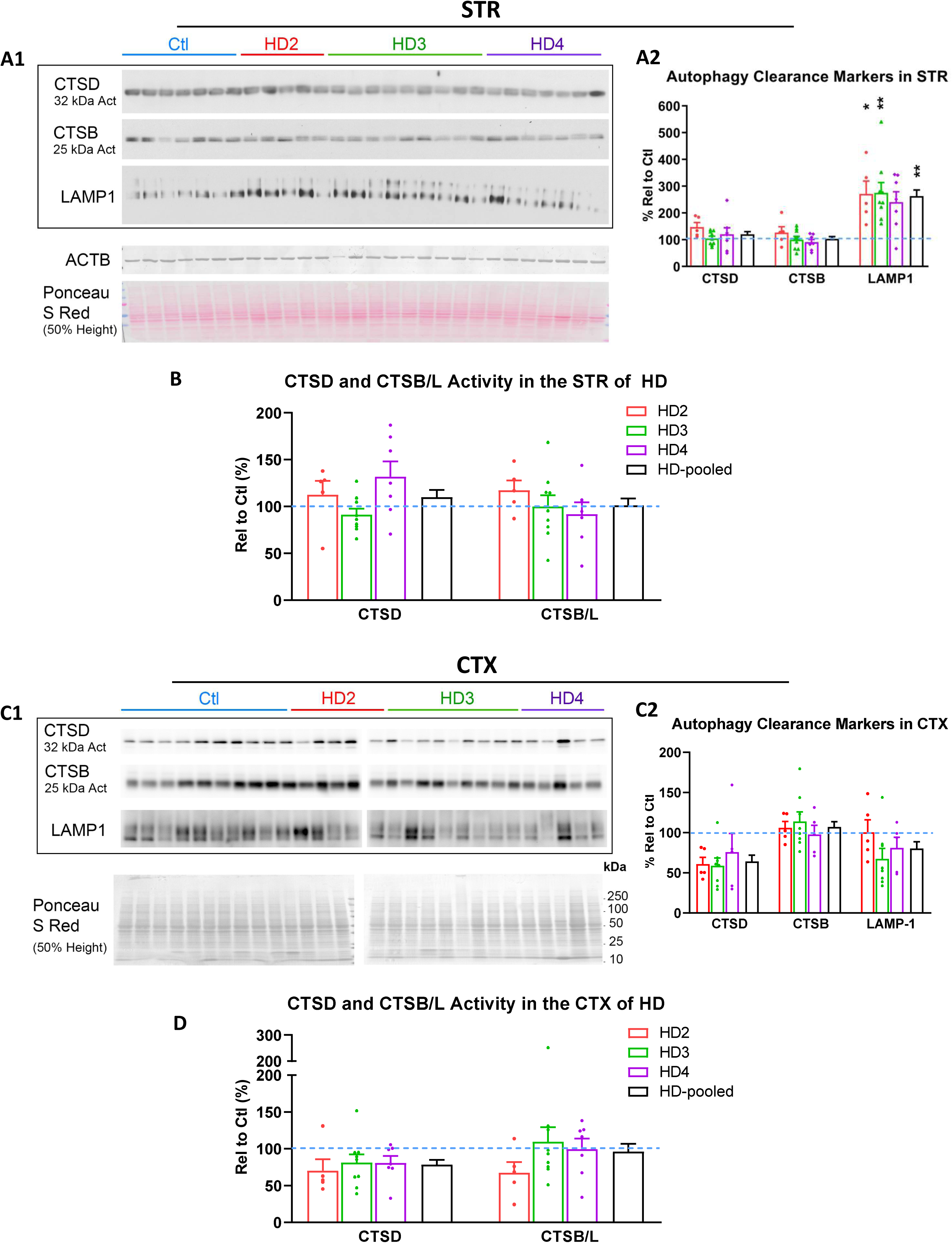
Levels of proteins involved in the lysosomal clearance phase of the ALP and enzymatic activity assays of cathepsins in the STR and the CTX. (A1 and C1) Representative western blots of striatal (A1) or cortical (C1) lysates demonstrating the levels of proteins involved in the lysosomal clearance stages of the ALP in control and HD cases. Color bars above the blots denote HD staging (Blue = Ctl; Red = HD2; Green = HD3; Purple = HD4). The western blots are accompanied with a representative colorimetric panel of ACTB as a loading control as well as a representative Ponceau S Red staining to highlight total proteins in samples shown (20 μg/lane). (A2 and C2) Bar graphs for quantitation results for the blots shown in (A1) and (C1), respectively. Each bar represents the result of either HD2, HD3, HD4 or the pooled data from all HD samples and is expressed as % of Ctl (set as 100% depicted by the dashed line) ± SEM. (B and D) Quantitation of CTSD and CTSB/L enzymatic activity in striatal (B) or cortical (D) lysates of controls or HD cases. Each bar represents the result of either HD2, HD3, HD4 or the pooled data from all HD samples and is expressed as % of Ctl (set as 100% depicted by the dashed line) ± SEM. Significant differences were analyzed by One-way ANOVA followed by post hoc Tukey’s multiple comparisons test. * signs: comparisons of each bar/group with the Ctl; # signs (if shown): comparisons among the groups. * P<0.05, ** P<0.01. For STR, n = 7 control and 21 HD cases; for CTX, n = 10 control and 18 HD cases.

We next examined transcripts of genes in the ALP that revealed a trend of upregulation of TFEB mRNA expression in the STR and CTX of HD and significant upregulation of LAMP1 and LAMP2 mRNAs in the STR of HD compared to control cases (Fig. 7), which may be the basis for the increased levels of LAMP1 protein in the STR of HD described above. No significant alterations were found in other transcripts including LC3, p62, CTSD, CTSB and HEXA in the STR and CTX of HD compared to control cases (Fig. 7).

**Fig. 7.**
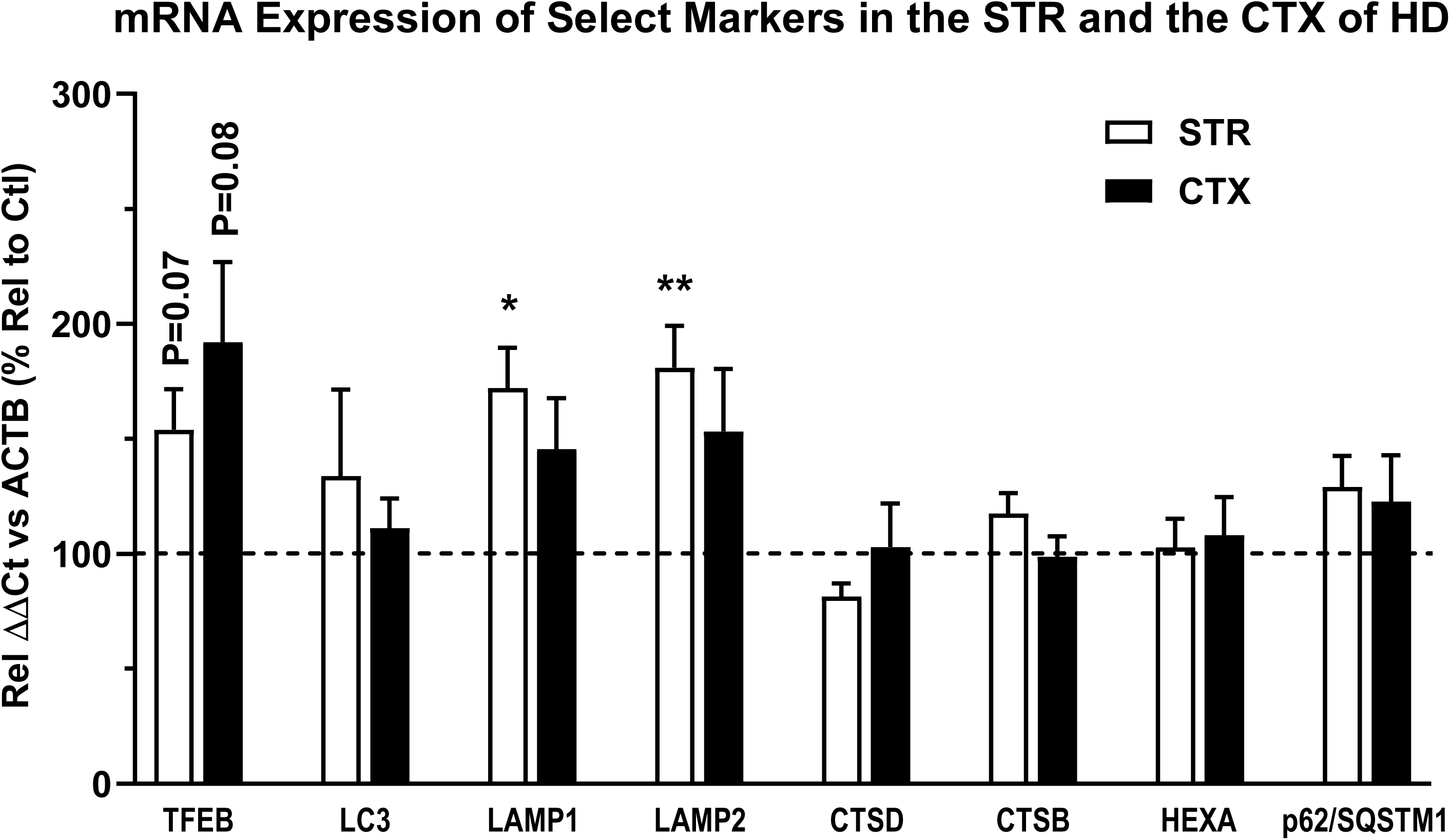
qPCR of selected ALP-related targets in HD brains. qPCR for analysis of mRNA transcripts from the STR and the CTX of control and HD cases (HD2 – HD4 stages) for assessing transcripts of genes involved in the ALP. Results are expressed as ΔΔCt relative to % of control ACTB levels ± SEM, with all HD staged levels pooled to yield a single result. Significant differences between the control and the HD case of each brain region, or between the STR and the CTX of HD case were analyzed by two-tailed Student’s t-test. * signs: comparisons with the Ctl; # signs (if shown): comparisons between STR and CTX. * or # P<0.05, ** or ## P<0.01, *** or ### P<0.001, **** or #### P<0.0001. For STR, n = 8 control and 20 HD cases; for CTX, n = 9 control and 21 HD cases.

## Discussion

Most of the knowledge about the relationship of the ALP with HD including the roles of HTT/mHTT in autophagy has been obtained from studies using cell and animal models. However, here, we analyzed a relatively large number of human brain samples from controls and HD2-HD4 patients, which has provided human brain-derived information that can greatly expand our knowledge about the status of ALP in HD brain, its relationship to the development of mHTT aggresome/inclusion pathology, and the relevance of animal models as surrogates to characterize HD neuropathology and pathobiology. Moreover, the findings we have described have implications as support for potential therapeutic value of specific strategies of autophagy modulation in HD.

### The status of ALP in the HD brain: STR vs CTX

At a global level, we have demonstrated in the human brain that HTT/mHTT are substrates of ALP. Moreover, in HD brain, our collective data suggest that ALP in affected neurons is relatively competent to maintain autophagy flux clearance capacity at an early disease stage, but that at later stages, autophagy flux declines and ALs laden with substrates including HTT accumulate. The later stage pattern is a pathological state associated with neurodegeneration in more than several major adult onset neurodegenerative diseases (Nixon and Rubinsztein 2024). A gradual bidirectional pathological relationship (a “vicious cycle”) is suggested between mHTT build-up within AL and the decline in autolysosomal clearance efficacy that is likely multifactorial (Nixon 2020, Nixon and Rubinsztein 2024). The relative competence of autophagy flux at an early disease stage of symptomatic HD contrasts with the temporal pattern in AD, where impaired flux emerges very early in pre-symptomatic disease as a result of direct impact of disease-related genetic factors on AL/LY proteolytic efficacy (Bordi, Berg et al. 2016, Nixon 2020, Im, Jiang et al. 2023, Nixon and Rubinsztein 2024) and autophagy failure progresses to an unusually extreme degree as disease advances (Lee, Yang et al. 2022)(Lee et al., 2024 abstract). Preservation of autophagy early in symptomatic HD suggests the potential opportunity to intervene therapeutically at this stage to stimulate autophagy flux with less concern than applies to AD where overburdening failing AL/LY and exacerbating build-up of toxic metabolites within the pathway may be counter-productive.

At the early HD2 stage, there is limited evidence suggesting a major alteration in autophagy induction and upstream steps of autophagy, such as AP formation or AP-LY fusion. We cannot, however, exclude a possible impairment in the engagement of certain autophagy cargoes by adaptor proteins which might impede sequestration and reduce ALP flux. We observed varying size clusters of double membrane limited vesicular profiles containing minimal intralumenal content (Fig. 5C1, C2), raising the possibility that they represent the “empty APs” described previously in mouse, cell, and patient fibroblast models of HD (Martinez-Vicente, Talloczy et al. 2010). Because these patterns were also often seen in fresh postmortem brains from individuals with AD clinical diagnoses in our other studies (Fig. S3A,B,D,E), but not seen in our previously published analyses of biopsied brain from AD patients (Nixon, Wegiel et al. 2005), we are inclined to attribute much of this pattern to postmortem and fixation artifact, mainly resulting in a swelling and vacuolization of endoplasmic reticulum. This is supported by the common observation in the HD brain of mitochondria within AP (mitophagy) and by our further immunogold labeling by an anti-calnexin antibody in HD brain which showed a significant decoration of these vesicular membranes (not shown). Artifactual vacuolization of lipid granules in lipofuscin may also give rise to a somewhat similar membrane pattern (Fig. S3C). Thus, our studies so far neither give significant support to the presence of empty APs nor refute the notion that they may be present in HD cells and cell models (Yang, Chen et al. 2021).

Interestingly, we detected spatial differences in the early ALP responses between the STR and the CTX. In the STR, but not the CTX, increased mRNA levels of LAMP1 and LAMP2 and protein levels of LAMP1 that imply lysosomal membrane expansion suggest a possible compensatory upregulation in lysosomal biogenesis. The absence of changes in levels of upstream ALP components (e.g., BECN1, ATG5, ATG7 and LC3) supports absence of deficiency in AP formation or compartment size although a caveat is the potential for glial cell autophagy systems to mask changes in neuronal levels in immunoblot analyses at the brain tissue level.

The less vulnerable but still affected CTX displayed no significant alterations in all the ALP markers examined biochemically, including enzymatic activities of lysosomal proteases and mRNA levels, suggesting that an apparently normal ALP machinery is maintained in this brain region. This absence of biochemical changes, together with the mild morphological changes [incl. AL clustering revealed by CTSD IHC (Fig. 3, top), inclusion formation revealed by mHTT IHC with mEM48 (Fig. S1c)], lower levels of aggregated HTT detected by immunoblotting with antibody MW8 (Fig. S2) and minimal HTT fragment generation (Fig. 2A2), highlight the regional difference of the CTX from the STR in the severity of pathological changes.

In neurons in both brain regions at late stages of disease, HTT/mHTT exists in CTSD-positive ALs which are abnormally enlarged and clustered. The continuing accumulation of these protein aggregates in the HD brain, especially in the STR, may be explained as a result of continuing overload of the aggregation-prone proteins on to the neurons which is beyond the degradative capacities of both macroautophagy (this study) and chaperone-mediated autophagy, which is known to play a role in HTT clearance and be upregulated in experimental HD models (Koga, Martinez-Vicente et al. 2011). Consistent with this, larger mHTT inclusions were observed in the neuropil without being completely contained within ALs (Fig. 4C), suggesting that increasing amounts of mHTT accumulation at the late stage of disease progression led to the formation of aggregates outside of vesicular structures that are too large to be processed by autophagy. However, we cannot exclude a possible mechanism that impairments in the interactions of adaptor proteins with autophagy cargoes could lead to slower rates of clearance of substrates including mHTT (Ochaba, Lukacsovich et al. 2014, Rui, Xu et al. 2015).

### The classification of inclusion bodies

We observed nuclear, neuritic and cytoplasmic inclusions and various subtypes in each category, particularly their ultrastructures. Although certain of these are described previously in literature (Tellez-Nagel, Johnson et al. 1974, Roizin 1979, DiFiglia, Sapp et al. 1997, Gutekunst, Li et al. 1999, Rudnicki, Pletnikova et al. 2008), they have not been presented collectively or described systematically and therefore what we present here may represent a relatively comprehensive collection of the inclusion types in the human HD brain. Among the NIIs, the main subtype is the pale-staining, spherical/ovoid fine granular and/or fibrous inclusion (Fig. 1A1, left). This may represent the most commonly reported inclusion type in brains of both human HD (Roizin 1979, DiFiglia, Sapp et al. 1997, Rudnicki, Pletnikova et al. 2008) and HD mouse models including R6/2, YAC128, HdhQ92 (Davies, Turmaine et al. 1997, Bayram-Weston, Jones et al. 2012, Bayram-Weston, Jones et al. 2012) and Q175 (our own study)(Stavrides, Goulbourne et al. 2024). Among the neuritic inclusion subtypes we identified (Fig. 1B), the major subtype (Fig. 1B3) also exhibited similar aggregate ultrastructure to the aforementioned main NII subtype (i.e., spherical/ovoid fine granular and/or fibrous inclusion), along with additional AVs and mitochondria inside and surrounding the inclusion. This subtype was also observed in other studies in the brain of human HD by HTT IEM (DiFiglia, Sapp et al. 1997, Gutekunst, Li et al. 1999) and HD mouse models (Bates, Mangiarini et al. 1998, Davies, Turmaine et al. 1999, Stavrides, Goulbourne et al. 2024). Again, similar aggregate ultrastructure of this type was also seen in some cytoplasmic inclusions (Fig. 1C1). Together, this fine granular and/or fibrous structure of aggregates appears to represent a major common ultrastructure, implying that the source of the aggregate material may be the same.

Another major ultrastructural feature of aggregates is a fiber-bundle subtype with variable shapes: rod- or comet-shaped or just parallelly arranged, which is more observed in the neuritic inclusions (Fig. 1B4, B5), but can be occasionally seen in the nucleus (Fig. 1A1, right, arrowhead) and the cytoplasm (Fig. 1A1, left, arrow). The fingerprint-like features of neuritic inclusion (Fig. 1B4) were considered as fibrillary fascicles from abnormal mitochondria in a previous study (Tellez-Nagel, Johnson et al. 1974). However, we tend to interpret them as HTT-derived fibrillar bundle aggregates as revealed from our high-resolution EM images (Fig. 1B4, enlarged images), and therefore include them within this fiber-bundle category.

These two ultrastructural aggregate types (compact fine granular and/or fine fibrous aggregate vs. fiber-bundle aggregate) apparently represent different ultrastructures, whereas they are very likely derived from and/or primarily composed of the same aggregating material, namely mHTT, particularly mHTT N-terminal fragments especially the Exon 1 fragments, which is supported by the following considerations. First, the direct evidence is that both types were positively labeled by anti-HTT antibodies by IEM in both human and mouse brains, and both fine granular/fibrous and bundle structures can be detected even in the same single inclusion (DiFiglia, Sapp et al. 1997, Davies, Turmaine et al. 1999, Gutekunst, Li et al. 1999). Second, studies using recombinant or overexpressed proteins of mHTT N-terminal fragments have demonstrated that in addition to oligomers and protofibrils, there were two main mature aggregation forms generated from the protein fragments, i.e., short fibrils and more aggregated bundles, which are similar to the two types of ultrastructures in the HD brain described above. Also, these studies found that the various patterns of aggregation existing during the aggregation process *in vitro* were interconvertible (Scherzinger, Lurz et al. 1997, Sahl, Weiss et al. 2012, Ko, Isas et al. 2018, Mario Isas, Pandey et al. 2021). However, it should also be noted that the aggregation formation process and the final ultrastructural morphologies of aggregates *in vitro* or *in vivo* can be affected by many factors such as experimental conditions, peptide sequence length, posttranslational modifications, and lipids, proteins and cellular membranes existing in an *in vivo* environment (Wang and Lashuel 2013, Kolla, Gopinath et al. 2021, Riguet, Mahul-Mellier et al. 2021).

### The significance of HTT and p62 fragmentation in HD

mHTT fragments are believed to be critical for the pathogenesis of HD and numerous studies have reported the presence of N-terminal and C-terminal fragments in human STR samples (Mende-Mueller, Toneff et al. 2001). Fragments may be generated by aberrant splicing of *HTT* or proteolytic cleavage of HTT (Sathasivam, Neueder et al. 2013). Particularly, more studies have focused on N-terminal fragments and revealed their pathological significance including their contribution to the formation of inclusions (Mangiarini, Sathasivam et al. 1996, Davies, Turmaine et al. 1997, Lunkes, Lindenberg et al. 2002, Schilling, Klevytska et al. 2007). Multiple sites for cleavages by proteases like caspases, calpains and for posttranslational modifications have been identified in the N-terminus (Kim, Yi et al. 2001, Gafni and Ellerby 2002, Wellington, Ellerby et al. 2002, Saudou and Humbert 2016, Sap and Reits 2020, Chiki, Zhang et al. 2021). In HTT-Knock-In mice, the majority of N-terminal fragments are most likely proteolytic products while the smallest fragment, i.e., the exon 1 protein, may be a product of incomplete splicing (Landles, Sathasivam et al. 2010).

An early study had reported the presence of HTT fragments of about 40 kDa in CTX samples from juvenile HD patients (65->70 CAG) but not in those from controls, and these fragments are the predominant species in the nuclear fraction (DiFiglia, Sapp et al. 1997), implying that they are the major species for NII formation. In our immunoblotting studies of the two brain regions, fragments of HTT (45-48 kDa) were not detected in the CTX but were readily detected in the STR where they were present at much higher levels in samples of HD patients than those of control cases. Together with the data from DiFiglia et al. (1997), our data suggest region-specific, disease severity-dependent, and/or CAG length related generation of the fragments. These 45-48 kDa species can be considered as N-terminal fragments as they were detected by the N-terminal antibody mEM48 but not by the Ab D7F7 which targets residues surrounding Pro1220 (not shown). Further, based on their size of 45-48 kDa, it is possible that they are generated from the cleavage at one of the calpain proteolytic sites, 437 (Gafni and Ellerby 2002), but not from other known calpain and caspase cleavages (i.e., calpain at 469 and 536; caspase at 513, 552 and 586) (Gafni and Ellerby 2002, Saudou and Humbert 2016) which would have generated larger fragments than 45-48 kDa. However, a possibility that these 45-48 kDa fragments can be the sequential proteolytic products from initial fragments generated by these proteases cannot be excluded (Kim, Yi et al. 2001, Gafni and Ellerby 2002). Moreover, suspected contribution of aspartic endopeptidases (Lunkes, Lindenberg et al. 2002) and/or matrix metalloproteinase (at aa 402) (Miller, Holcomb et al. 2010) to the generation of these fragments may also be considered. On the other hand, such interpretations may not be accurate or necessary given that many posttranslational modifications may occurs at the N-terminus (Saudou and Humbert 2016) and that the gel migration of the HTT fragments are retarded by the expanded polyQ tract as mentioned previously (Sathasivam, Neueder et al. 2013, Landles, Milton et al. 2020). All of these possibilities for fragment speciation make it difficult to establish the actual aa sequence size and the responsible protease(s). However, no matter how these fragments are generated, their specific increase in the HD STR (vs the controls) starting at HD2 may suggest their involvement in inclusion formation and proclivity to form inclusions rather than indicate only a deficit in their clearance by the proteolytic systems.

In a number of HD mouse models, exon 1 protein generated as a consequence of aberrant splicing can be detected by MW8, an antibody targeting the C-terminus of exon 1 protein and capable of detecting the aggregated form of the exon 1 HTT protein, where the size of the exon 1 protein on the gel varies depending on the size of the polyQ (Sathasivam, Neueder et al. 2013, Landles, Milton et al. 2020). However, this exon 1 protein was not detectable with MW8 in cerebellum (CBM) homogenates from HD patients (Neueder, Landles et al. 2017). In the current study, we were interested in examining if exon 1 protein was present in STR and CTX samples with MW8. Although we could detect aggregated HTT in the stacking gel in samples from STR (Fig. 2A1, Row 1; Fig. S2) and barely in a few cases of CTX samples (Fig. 2A2, Row 1; Fig. S2), we failed to detect any bands below 250 kDa as shown on the full-size blots (Fig. S2). This result is not surprising given that the levels of exon 1 mRNA in CTX, CBM and hippocampus are only significantly higher in juvenile HD cases but are very low in adult-onset HD patients, which are not significantly different from those of controls (Neueder, Landles et al. 2017). It is notable that more recent studies in human brains using a highly sensitive approach detected fair normalized HTT1a (to HTTex2) expression levels in samples with CAG sizes in the juvenile-onset range. However, the two samples with repeats in the adult-onset range only show very low HTT1a/HTTex2 ratio, similar to the levels in the controls (Hoschek, Natan et al. 2024). Similarly, aberrant spliced HTT exon1 could be detected in the brains of HD pigs, but it was expressed at a much lower level than the normally spliced HTT exon products (Tong, Yang et al. 2023). Together, even though it cannot be ruled out at this point that our immunoblotting approach is not sensitive enough to detect the very levels of exon 1 protein, if any, the absence of MW8 band(s) below 250 kDa in our study, including the 45-48 kDa range, suggests that the 45-48 kDa fragments in our STR samples (Fig. 2A) are not likely to be the exon 1 protein, but are proteolytic products discussed above. Further studies may be required to verify if exon 1 protein is actually produced in human brain samples, like in the brain of mouse models. A much higher level of aggregated HTT in the stacking gel in HD STR samples compared to HD CTX samples is striking, suggesting many more aggregates (in the form of inclusions, and maybe in other forms as well) exist in the STR which would be apparently consistent with a general view that the STR is a more affected region than the CTX. These differences in the levels of the aggregated HTT (by MW8) and the 45-48 kDa HTT fragments (by mEM48) between the STR and the CTX may be an underlying mechanism for the different degrees of lysosomal membrane expansion/AL accumulation between these brain regions discussed above.

We also observed increased levels of p62 fragments in the STR samples of HD compared to controls. Previous *in vitro* studies have found that caspases cleave p62 at several putative sites (D256, D329, D337 and D347), generating fragments of various sizes (30, 35, 37, 40, 46 kDa) (Norman, Cohen et al. 2010, Jamilloux, Lagrange et al. 2018, Sanchez-Garrido, Sancho-Shimizu et al. 2018, Valionyte, Yang et al. 2022). Among these cleavages, cleavage at D329 by caspase-1 generates a 30 kDa fragment and diminishes p62-LC3 interaction (Jamilloux, Lagrange et al. 2018), or by caspase-8 generates a 40 kDa fragment and promotes mTOR activity (Sanchez-Garrido, Sancho-Shimizu et al. 2018); and cleavage at D256 by caspase-6 produces a 30 kDa fragment and attenuates p62-droplet dependent AP formation (Valionyte, Yang et al. 2022), all indicating negative effects on autophagy. In our present *in vivo* study, while we detected fragments with similar sizes (ranging 32-48 kDa), we are not sure if any of the fragments correspond to any found *in vitro*. Particularly, the anti-p62 antibody we used is directed against the C-terminal 20 aa, and therefore we would have seen C-terminal fragments of ∼ 20 kDa or smaller if caspase-medicated cleavages did occur (i.e., the largest C-terminal fragment released from the 440-aa full-length p62 would be ∼184 aa if cleaved at D256 by caspase-6). Thus, the fragment size range of 32-48 kDa implies that other proteolytic events rather than/in addition to the caspase cleavages may be involved. A potential contribution of calpain in the generation of the observed fragments remains to be studied even if an *in vitro* study did not detect fragments after treating p62 with calpain 1 which suggested that calpain 1 is capable of completely digesting p62 *in vitro*, without leaving detectable fragments (Norman, Cohen et al. 2010). Nevertheless, in terms of negative effects on autophagy under a caspase-cleavage situation, our data, which indicate a moderate compensatory upregulation in lysosomal biogenesis without signs of blockage in autophagy steps (e.g., AP formation or AP-LY fusion) as mentioned above, do not support an existing autophagy inhibition as an explanation for the p62 fragments. That a high level of fragments is present at HD2 when the LY function is intact does not support a likelihood that increased levels of p62 fragments are due to an impaired clearance of these fragments in the ALP.

In summary, our main observations of ALP alterations in the STR from this study include: moderately upregulated lysosomal biogenesis reflected by increased LAMP1 and LAMP2 marker levels, engagement of mHTT as an ALP substrate, absent upstream autophagy pathway deficits, and emergence of AL enlargement/clustering at late disease stages (mainly in HD4). These findings together suggest that the autophagy machinery and the rates of autophagy flux in HD brain may not significantly decline until late when mHTT degradation becomes delayed. A striatum specific finding is the high levels of fragments of HTT, p62 and TRAF6, three cross-interacting proteins (Zucchelli, Marcuzzi et al. 2011, Linares, Duran et al. 2013), appearing at HD2 and persisting to HD4, which may primarily reflect enzymatic cleavages promoting formation of inclusions rather than defective lysosomal clearance. The observations in HD revealing mild AV accumulation and AL enlargement even at HD4 along with preserved lysosomal enzymatic activities contrast with an AD-like early and progressive lysosomal proteolytic decline and massive autophagic substrate build-up. These contrasting situations suggest that, unlike concerns for AD therapy (Nixon and Rubinsztein 2024), pharmacologic stimulation of autophagy in HD beginning in early symptomatic stage, as proposed by others (Ravikumar, Vacher et al. 2004, Sarkar and Rubinsztein 2008), may well show greater efficacy in eliminating autophagic substrates including mHTT in HD while avoiding ALP flux backup due to lysosomal degradation disruption. This idea has been successfully verified in our study in the Q175 mouse model (Stavrides, Goulbourne et al. 2024) where administration of a mTOR inhibitor to 6-mo-old Q175 normalized LY number, ameliorated aggresome pathology while reducing HTT-, p62- and Ub-IR, suggesting beneficial potential of autophagy modulation at early stages of disease progression.

## Acknowledgement

This work was supported by the CHDI Foundation (R.A.N.) and the National Institute of Aging (P01 AG017617 to R.A.N.). We are very grateful to all brain banks for providing the valuable human brain samples, including Harvard Brain Tissue Resource Center, Emory Center for Neurodegenerative Disease, New York Brain Bank at Columbia and Mount Sinai Neuropathology Brain Bank & Research Core.

## Materials and Methods

### Brain tissue

Brain samples were obtained from the following brain banks: Harvard Brain Tissue Resource Center (HBTRC), Emory Center for Neurodegenerative Disease (ECND) and New York Brain Bank at Columbia (NYBBC). These banks use Vonsattel’s grading system of neuropathological severity to stage brains from individuals diagnosed clinically as having HD as Grade 0 to Grade 4 (HD0 – HD4) (Vonsattel, Myers et al. 1985). Two brain regions were used in this study (STR = caudate nucleus of the striatum, CTX = prefrontal cortex) as indicated in **Table 1** which provides detailed demographic information.

### Antibodies for immunohistochemistry (IHC), western blotting (WB)

The following primary antibodies were used in this study. (1) from Cell Signaling Technology: tHTT rabbit mAb (clone D7F7, #5656, targeting residues surrounding Pro1220 of human HTT and detecting total HTTs), p70S6K pAb (#9202), p-p70S6K (T389) pAb (#9205), ULK1 pAb (#4773), p-ULK1 (S757) pAb (#6888, #14202; detecting S757 or S758 of mouse or human ULK1, respectively), ATG5 rabbit mAb (#12994), ATG7 pAb (#2631), ATG13 rabbit mAb (#13273), p-ATG13 (S355) rabbit mAb (#26839), VPS34 rabbit mAb (#81453), TRAF6 rabbit mAb (#8028). (2) from Millipore-Sigma: ntHTT mAb (N-Terminus-specific, mEM48, #MAB5374, preferentially recognizing aggregated HTT)(Gutekunst, Li et al. 1999), ATG5 pAb (#ABC14), K48- or K63-specific ubiquitin mAb (#05-1307, #05-1308, respectively), βIII-tubulin mAb (#SAB4700544), β−actin mAb (#A1978). (3) from other vendors: BECN1 mAb (BD Biosciences, #612113); LC3 pAb (Novus Biologics, #NB100-2220), ATG9 (Novus Biologics, #B-110-56893); p62 mAb (BD Biosciences, #610832) or C-term-specific p62 Guinea Pig pAb (Progen Biotechnik #C-1620); total ubiquitin pAb (Dako Agilent, #Z0458), LAMP1 or LAMP2 rat mAb (Developmental Studies Hybridoma Bank, University of Iowa, #H4A3 or #H4B4, respectively); CTSD sheet pAb (D-2-3, in-house made) (Cataldo, Thayer et al. 1990); CTSD pAb (Scripps Laboratories, #RC245), CTSD mAb (CD1.1, in-house made) (Lie, Yang et al. 2021); CTSB pAb (Cortex Biochemicals, #CR6009RP), CTSB goat pAb (Neuromics, #GT15047).

The following secondary antibodies and reagents for immunoperoxidase labeling were purchased from Vector Laboratories (Burlingame, CA): biotinylated goat anti-rabbit or -mouse IgG/IgM, Vectastain ABC kit (PK-4000), and DAB Peroxidase Substrate Kit (SK-4100). The following secondary antibodies for immunofluorescence were purchased from Thermo Fisher Scientific (Waltham, MA): Alexa Fluor 568-conjugated goat anti-mouse IgG (A11031), Alexa Fluor 488-conjugated goat anti-rabbit IgG (A11034), and Alexa Fluor 568-conjugated goat anti-rabbit IgG (A11036).

### Immunolabeling of brain sections

Formalin-fixed, tissue blocks of human brain (**Table 1**) were sectioned at 40 μm on a vibratome, or paraffin embedded and sectioned at 7 μm. Sections were deparaffinized as necessary. Antigen-retrieval was performed by boiling sections in sodium citrate buffer at 95°C for 30 minutes. Sections were blocked and incubated in primary antibody O/N (up to 3 days in some cases) at 4°C. Alexa-Fluor conjugated secondary antibodies were used for immunofluorescence and ABC detection method was used for immunoperoxidase labeling with DAB. Autofluorescence was quenched with 1% Sudan black (Sigma-Aldrich; St. Louis, MO) in 70% ethanol for 20 minutes. DAB labeling was inspected on a Zeiss AxioSkop II equipped with a HrM digital camera (Carl Zeiss, Germany). Immunofluoescent images were collected on a Zeiss LSM510 Metal confocal microscope.

### Ultrastructural analyses

For EM, vibratome sections of human brain (**Table 1**) were post-fixed in 1% osmium tetroxide. Following alcohol dehydration, sections were embedded in Epon (EMS, Hatfield, PA). One-micron-thick sections were stained with toluidine blue for light microscopic examination and ultrathin sections prepared and stained with uranyl acetate and lead citrate. Material was viewed with a Philips CM 10 electron microscope equipped with a digital camera (Hamamatsu, model C4742-95) aided by AMT Image Capture Engine software (version 5.42.443a).

Post-embedding IEM with gold-conjugated secondary antibody was performed to detect mHTT (antibody mEM48), pan-Ub, p62 and CTSD signal in neuronal cell bodies and the neuropil using a previously described protocol (88). Ultrathin sections were placed on nickel grids, air-dried, and etched briefly with 1% sodium metaperiodate in PBS followed by washing in filtered double-distilled water and incubated with 1% BSA for 2 hours. Sections then were incubated overnight in the anti-CTSD antibody (RU2, 1:1000) in a humidified chamber overnight at 4°C, washed in PBS, and incubated in a secondary antibody conjugated with 10-nm gold particles (Amersham, Buckinghamshire, UK) for 2 hours at room temperature. Grids were washed and briefly stained with uranyl acetate and lead citrate before examination.

Please note that the majority of EM images were from one HD4 case which had shorter PMI and exhibited an exceptionally high level of preservation of ultrastructure compared to over 20 postmortem HD brains surveyed. However, we did examine additional HD4 and HD3 cases (**Table 1**) with sufficient preservation of features to provide similar info to what we found from the above HD4 case, although their ultrastructure were suboptimal for performing quantitative EM analyses.

### Preparation of tissue extracts

Grey matter (0.5 g) was dissected from the STR and the CTX (Brodmann’s area 9/10) of human brains (**Table 1**) and homogenized in RIPA buffer (50 mM Tris/HCl pH 7.4; 0.15 M NaCl; 5 mM EDTA; 1 mM EGTA; 0.5% Sodium deoxycholate; 1% NP40, 0.1% SDS with protease inhibitors 1 mM AEBSF (Gold Biotechnology, St. Louis, MO) and 20 μg/ml of leupeptin and pepstatin (US Biochemicals, Cleveland, OH), and phosphatase inhibitor microcystin LR (1 ng/ml, Enzo). Lysates were frozen and thawed 3 times followed by centrifugation at 10,000 g for 30 min to yield a total tissue lysate supernatant. Protein content was determined by the BCA method (Smith, Krohn et al. 1985). Brain lysates were examined by western blotting for various marker proteins for the ALP.

### SDS-PAGE and western blotting

Lysate extracts (10-40 μg total protein) were separated on 4-20% or 10% Tris-glycine SDS-PAGE gels and transferred to nitrocellulose (Pall, Pensacola, FL) for probing with antisera as noted along with appropriate external controls. Blots were blocked for 1 hr at 37°C in 1x TBST containing 5% blotting grade dry milk (W/V), incubated in 1° Ab in block solution O/N at 4°C, washed 3x 10 min in TBST at RT followed by incubation with 2° Ab conjugated to horseradish peroxidase diluted in block solution for 90 min at RT. Blots were washed 3x 10 min and immunoreactive bands were visualized with ECL reagent (RPN2209, Millipore Sigma) and the bands were quantified using MultiGauge V. 3.0 (Fuji Film) software. Target proteins were normalized against β-actin, unless otherwise noted.

Exposures were approximated for multiple conditions based on levels found for external controls as noted in Results. In some cases, especially in cases of assaying housekeeping or other highly expressed antigens that exposure times are difficult to control and maintain signal linearity on film, blots were developed using DAB enhanced with nickel. Blots were stripped in 0.2 M HCl, pH 2.2 containing 0.1% SDS and 1% Triton X-100 for 1.5 hr at 25°C and reprobed using the above protocol.

### RNA preparation

For routine qPCR analyses 100 mg of grey matter from the STR and the CTX (Brodmann’s area 9/10) of human brains (**Table 1**) was dissected and extracted in 1.5 ml Trizol reagent (ThermoFisher/Life Technologies, 15596026) using a hand-held homogenizer (6 x15 s bursts)(Pro-Scientific, Oxford, CT) followed by mixing with 300 μL of chloroform. Samples were centrifuged at 12000 g for 15 min at 4 °C. The aqueous phase was collected and 750 μL isopropanol was added; samples were spun again at 12000 g for 10 min at 4°C. Supernatant was removed and pellet washed 2 times with 75% ice-cold ethanol and centrifuged 7500 g for 5 min at 4°C. The pellet was redissolved in 100 μL of RNAase-free distilled water. Total RNA quality (RNA Integrity Number, RIN) was assessed on an Agilent Bioanalyzer using an RNA Nanochip (2100; Agilent Technologies; Santa Clara, CA). RNA quantity was interpolated from the Agilent chip by using an RNA ladder with a known concentration of 150 ng.

### Preparation of cDNA and qPCR

Starting concentrations of total RNA were normalized for samples whose concentrations were estimated on different Agilent chips. cDNA was prepared using TaqMan Reverse Transciption Reagent kit (Applied Biosystems; Branchburg, NJ) according to manufacturer’s instructions. Following reverse transcription, sample cDNA was loaded in triplicate into wells of a 96-well optical reaction plate containing appropriate target gene primer (Applied Biosystems, Branchburg, NJ). GAPDH (glyceraldehyde 3-phosphate dehydrogenase), ACTB (actin β), and HPRT1 (hypoxanthine phosphoribosyltransferase 1) were run as housekeeping genes, also in triplicate, for each sample and on the same plate, as endogenous controls. Total reaction volume per well was 20μL. qPCR was performed in the ABI Prism 7900HT Sequence Detection System (Applied Biosystems Branchburg, NJ) as described previously (Alldred, 2015; Alldred, 2015).

### Calculation of qPCR results

Following qPCR, the target genes were normalized against the three housekeeping genes (GAPDH, ACTB, and HPRT). Results were calculated using the ΔΔCt method (Applied Biosystems, Branchburg, NJ Bulletin #2). Control values were averaged as a geometric mean, and sample values were recalculated and expressed as percent control. Outliers were recognized as values falling beyond two standard deviations of mean, and were discarded from the analyses.

### Enzymatic assays in brain homogenates

Cathepsins B and L were assayed by measuring the release of 7-amino-4-methylcoumarin (amc) from Z-Phe-Arg-amc at pH 5.5 (substrate recognized by both enzymes (Enzo, Plymouth Reading, PA) modified from the method of Barrett and Kirschke (Barrett, 1981) to utilize microplate procedures. Typically, assays were performed in black microplates in a volume of 100 μl mixture (1-5 μl of enzyme in 50 mM NA-Acetate, pH 6.5 containing 1 mM EDTA and 10 μM Z-Phe-Arg-amc). Fluorescence of amc released was read at different time points in a Wallac Victor-2 spectrofluorimetric plate reader with a filter set optimized for detection of 4-methyl-7-aminocoumarin (-*amc*) standard solution with excitation at 365 nm and emission at 440 nm. The reaction was linear up to 2 hours. Enzyme activity was expressed as the amount of amc released per hour per mg protein.

Cathepsin D was assayed at 37°C at pH 4.0 by measuring the release of amc containing peptide, 7-methoxycoumarin-4-acetyl-Gly-Lys-Pro-Ile-Leu-Phe from 7-methoxycoumarin-4-acetyl-Gly-Lys-Pro-Ile-Leu-Phe-Phe-Arg-Leu-Lys (Dnp)-D-Arg-NH2 (BioMol-Enzo, Plymouth Reading, PA), according to the method of Yasuda et.al. (Yasuda, 1999). Assays were performed in black microplates in a total volume of 100 μl (0.1 M sodium acetate buffer pH 4.0 containing 20 μM substrate with and without 3 μg of pepstatin) for one hour. Fluorescence released was read in a Wallac Victor-2 Spectroflurimetric plate reader with a filter optimized for detection of amc standard solution with excitation at 365 nm and emission at 440 nm. However instead of using amc standard, a quenched standard 7-methoxycoumarin-4-acetyl-Pro-Leu-OH was used for expressing enzyme activity to account for the release of peptide containing amc instead of free amc. Enzyme activity was expressed as the relative amount of quenched standard released per hour per mg protein. The specific activity of cathepsins were calculated by calculating the ratio of enzyme activity to the densitometric data obtained from western blots for each enzyme.

## SUPPLEMENTAL FIGURE LEGENDS

**Fig. S1 (related to Fig. 1).**
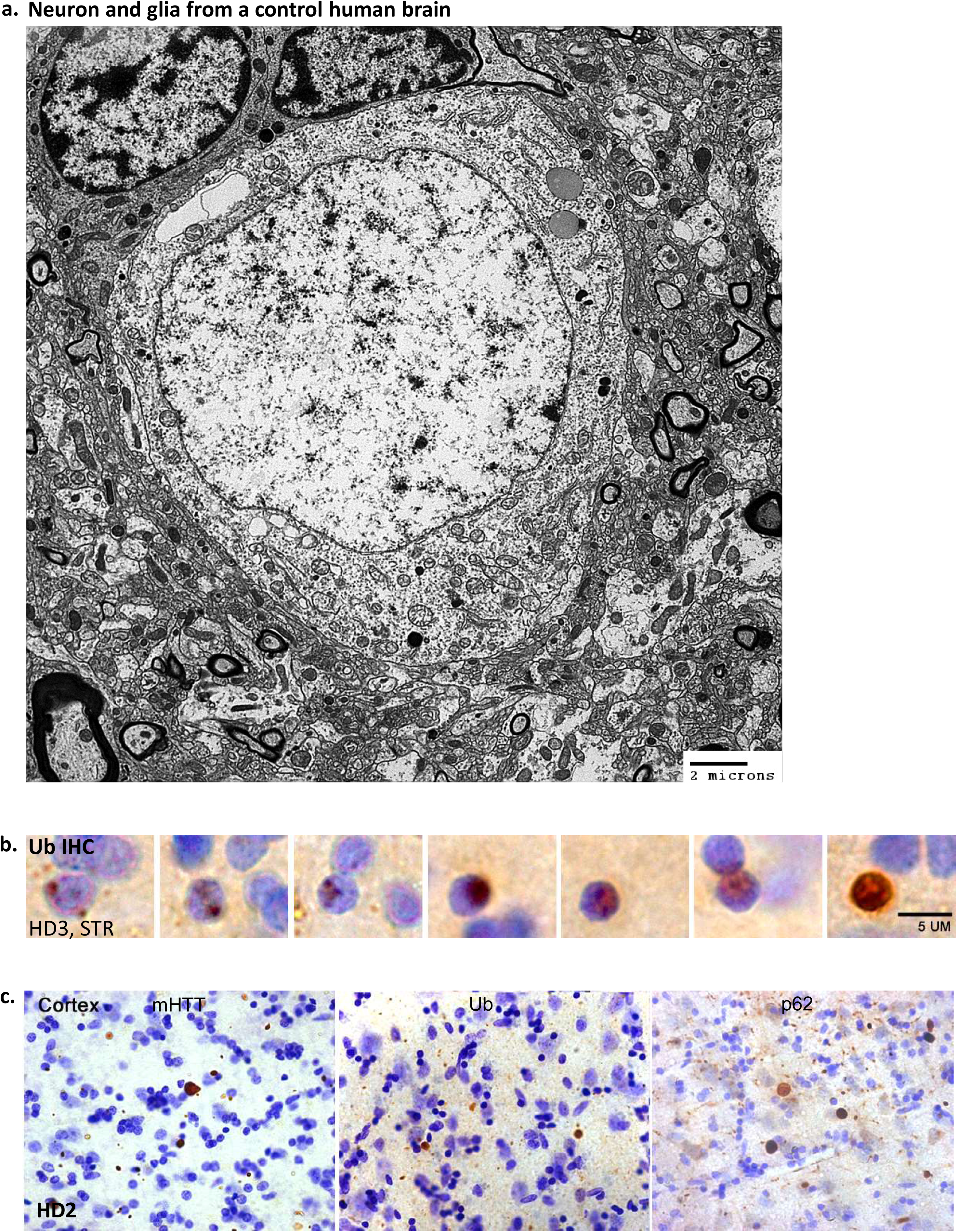
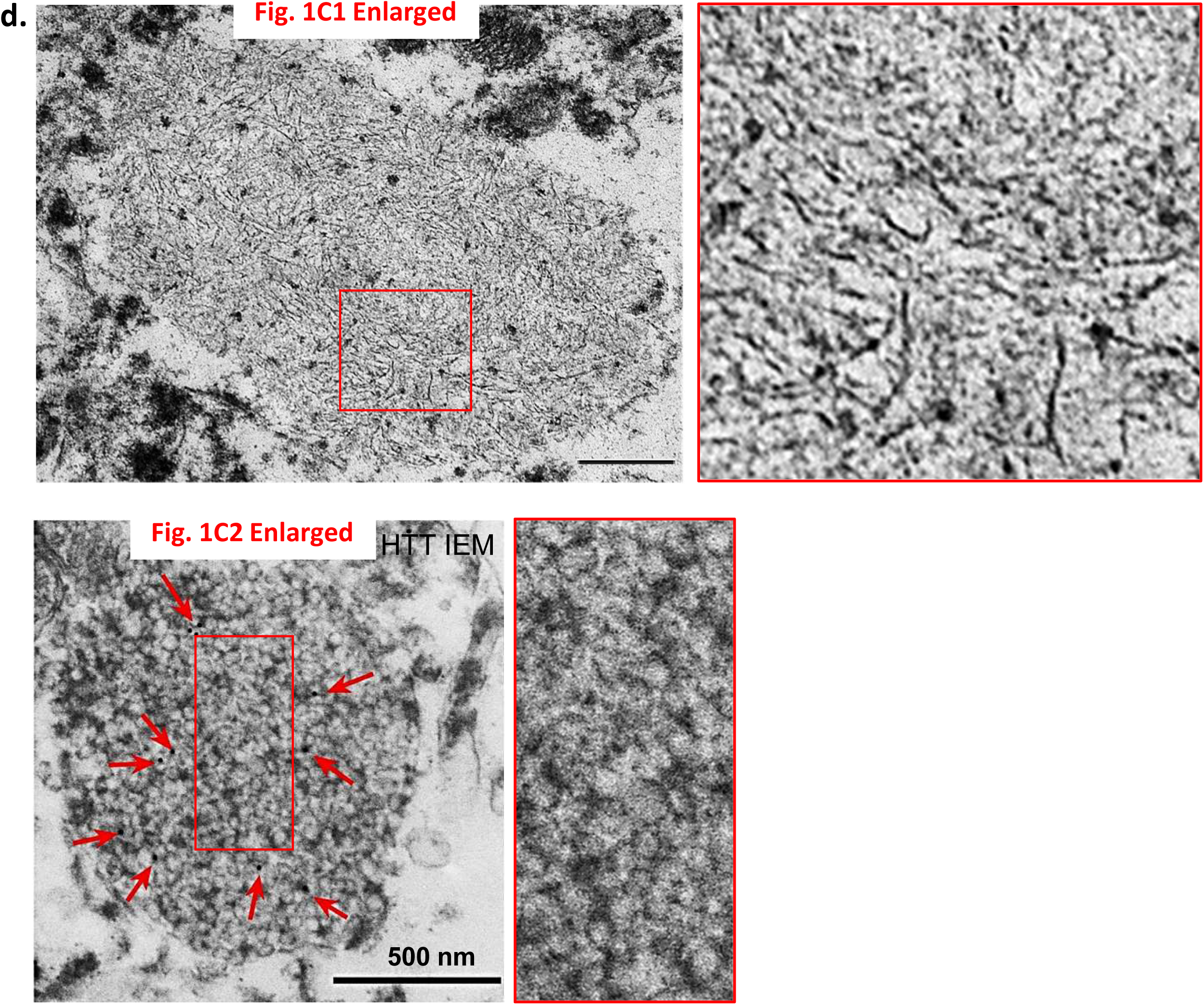
**a.** A representative EM image from a control human brain demonstrating a neuron (center) and two glial cells (top), as a reference for normal ultrastructure of neuronal perikarya. **b.** A series of LM images from the STR at the HD3 stage depicting the varying sizes of the Ub- positive NIIs. **c.** LM images showing neuritic inclusions positive for either mHTT (mEM48), pan-Ub or p62 randomly distributing in the neurpil in the CTX, similar to the pattern in the STR shown in main Fig. 1. **d.** These are the regular size (i.e., uncompressed) EM images shown in main Fig. 1C, facilitating visualization of detailed ultrastructures.

**Fig S2 (related to Fig. 2).**
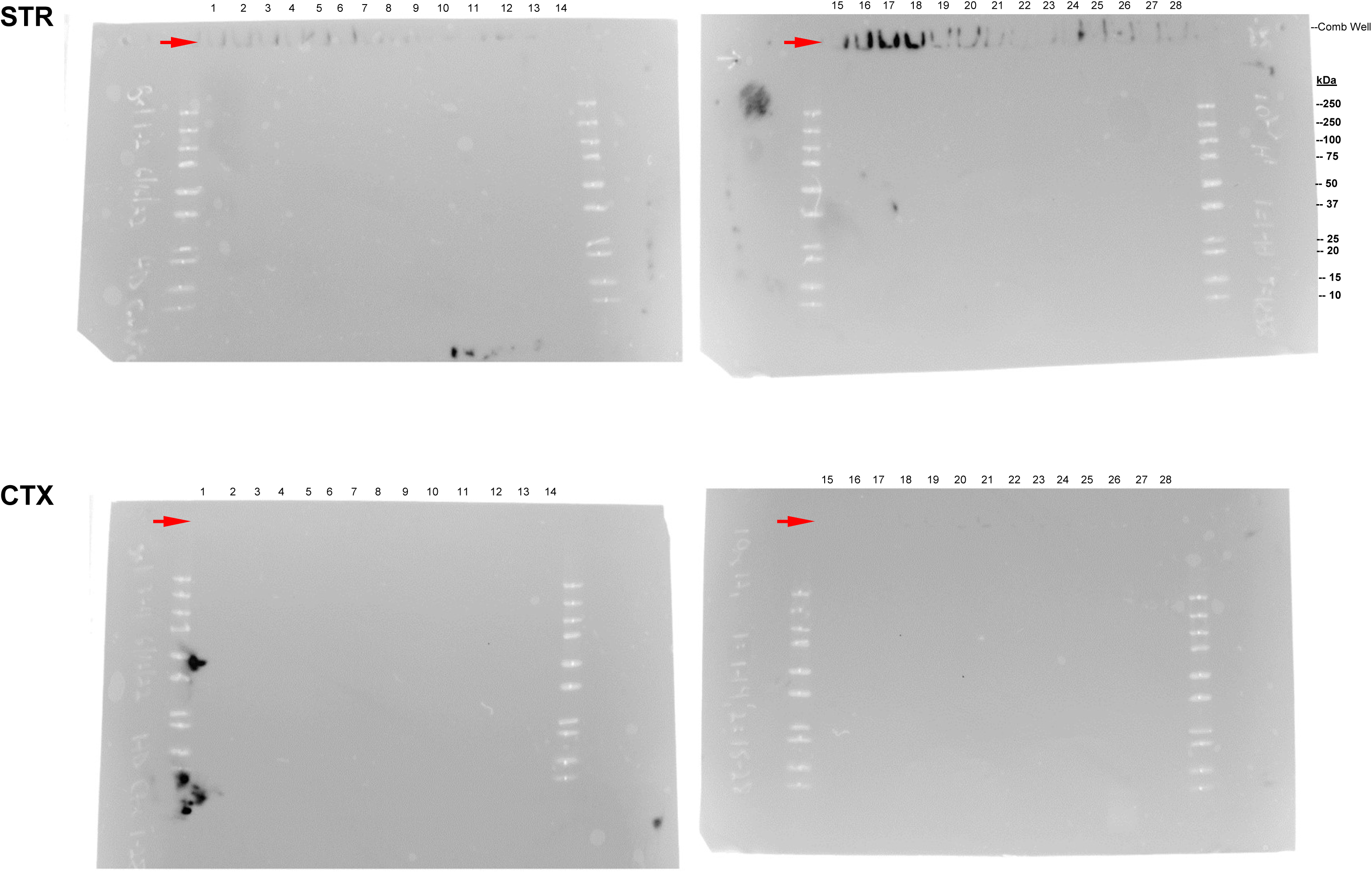
mHTT signal is only detected in the stacking gel but not below by antibody MW8. These are the full-size blots of those MW8 strips shown in Fig. 2 (A1, A2, Rows 1). Brain homogenates from STR and CTX of HD patients and controls were probed with mAb MW8 (Refer to the above Fig. 2 Legend for additional info). Shown are overlayed images of chemiluminescence signal representing the immunostaining and visible light (for easy viewing Mr Markers). Red arrows depict mHTT aggregates detected in the stacking gels which have been shown in Fig. 2A1, A2, Rows 1. No additional immunoreactive species are detected below the 250 kDa marker.

**Fig S3 (related to Fig. 5C).**
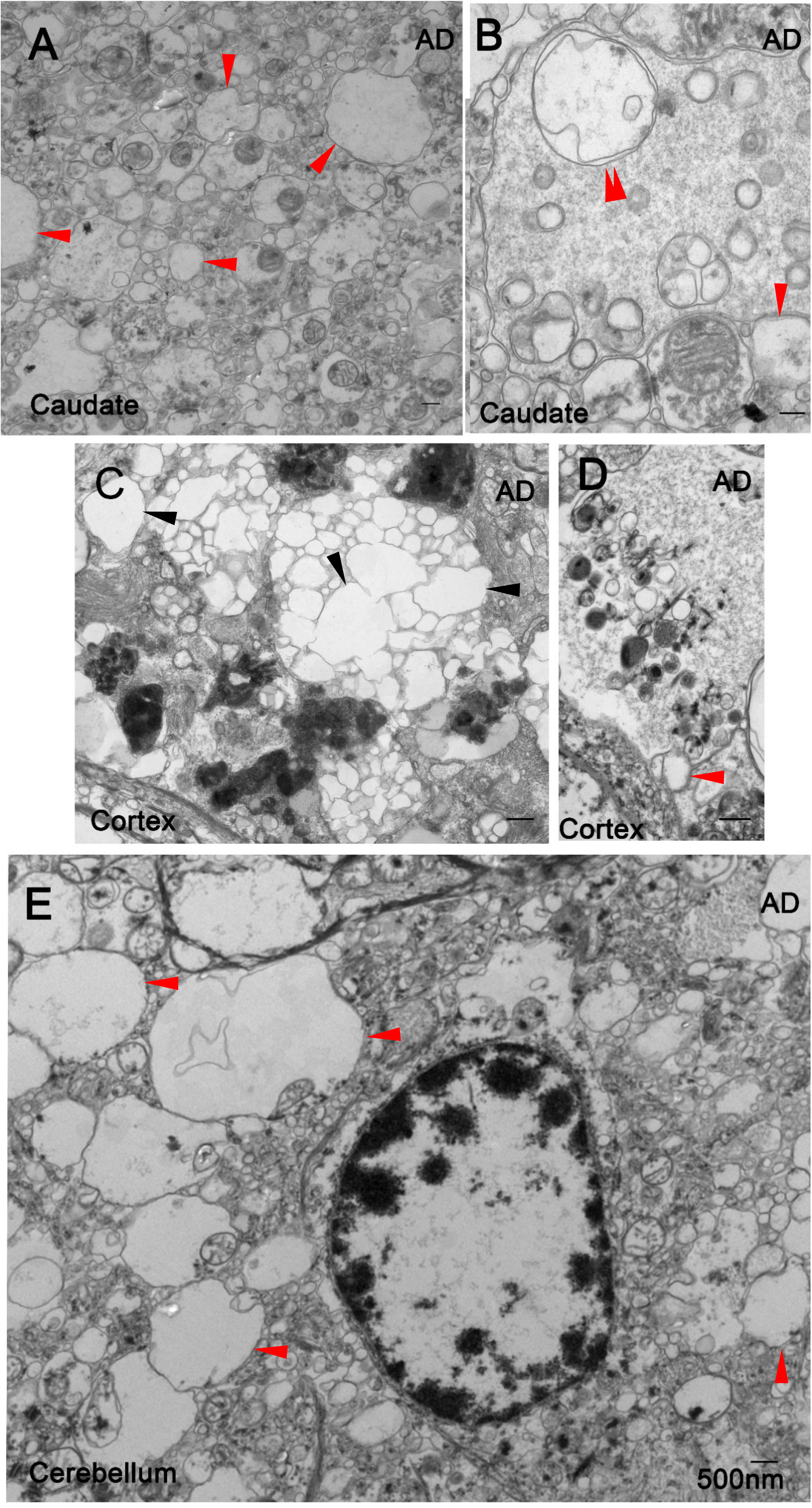
Membrane pathology observed in the AD brain. EM images collected from different brain regions of individuals with AD clinical diagnoses, demonstrating single or double membrane-limited clear/empty vesicles of varying sizes with minimal intralumenal contents (A, B, D, E, red single or double arrowheads, respectively), or artifactual vacuolization of lipid granules within lipofuscin granules (C, black arrowheads). Bars = 500 nm.

